# Genomic status of the Eurasian curlew *Numenius arquata*: estimating Essential Biodiversity Variables and selection signals for a declining migratory bird

**DOI:** 10.64898/2026.08.25.746821

**Authors:** Grace Walsh, Jacob Höglund, Patrik Rödin-Mörch, James A. Ward, Rebecca C. Örnberg, Jamie Thompson, Declan O’Donovan, Adriaan de Jong, Seán B.A. Kelly, Nicola Hemmings, David E. MacHugh, Barry J. McMahon

## Abstract

Understanding how contemporary population declines affect the genomic diversity and structure of threatened species is important for effective conservation. The Eurasian curlew (*Numenius arquata*) is experiencing severe population declines across Europe, with Ireland among the most extreme, showing declines exceeding 90% over 40 years. Genomic data are increasingly incorporated into policy and used to assess conservation status by estimating genetic diversity, differentiation, inbreeding, effective population size, and adaptive divergence. Such data for curlew is scarce, and the population structure among northern and north-western European breeding populations remains unclear. To address this, we generated whole-genome resequencing data for 56 curlews across Ireland, Britain and Sweden. Irish and British populations showed minimal interpopulation differentiation, but both were substantially differentiated from Sweden. This was apparent from principal component analysis, and admixture and *F*_ST_ analyses. Measures of genetic diversity (nucleotide diversity, heterozygosity, Watterson’s *θ*) were similar across populations. A slightly elevated Tajima’s *D* in Ireland, along with elevated *F*_ROH_ in Ireland and Britain relative to Sweden, may be the early genomic signs of recent population declines. We identified locally selected candidate genes. These had putative roles in metabolic processes, the immune response, and were potentially associated with distinct migratory behaviours and environmental conditions. We find a potential lag in genomic effects of decline being detectable following population contraction. We also show highly migratory species can exhibit differentiation in ecologically relevant traits, potentially driven by high site fidelity. These findings warrant consideration in translocation planning and broader conservation strategies.

## Introduction

Nature provides many services to people, such as biological pest control, nutrient transport, and seed dispersal, along with its inherent cultural value (Díaz-Siefer et al. 2022; Gaston et al. 2018). However, nature’s power to provide these services is increasingly compromised. As habitats fragment and populations decline, gene flow is reduced. The resulting small populations lose genetic diversity quickly due to genetic drift. This process can overpower the beneficial effects of natural selection (Allendorf et al. 2022), impacting population viability and adaptive capacity (Frankham 2005). Small populations also experience greater inbreeding, limiting population growth and fitness (Kardos et al. 2023) potentially leading to an “extinction vortex” where loss of genetic diversity through drift and inbreeding depression further reduces population size (Fagan and Holmes 2006). Over generations, this leads to genomic erosion where diversity is reduced, deleterious alleles segregate, and local adaptations can be lost (Bosse and Van Loon 2022). In addition, historically large populations typically have more genetic diversity, and therefore more deleterious variation, which could be expressed through inbreeding after a bottleneck (Beichman et al. 2023; Kyriazis et al. 2021).

Studying the overall genomic make-up of species can inform the conservation of multiple genetically distinct populations, which is required to support species survival and stable ecosystem functioning (Des Roches et al. 2021; Heuertz et al. 2023). Standardised metrics are needed to monitor changes in genetic diversity over time, such as the Essential Biodiversity Variables (EBVs) (Hoban et al. 2022): genetic diversity, genetic differentiation, effective population size, and inbreeding. EBVs provide a baseline for assessing the effectiveness of conservation efforts and population trajectories, aiding management and monitoring programme design, and translating well to policy. Genetic and genomic data have begun to enter policy, such as the Kunming–Montreal Global Biodiversity Framework (KMGBF) (Da Silva et al. 2025; Hoban et al. 2022). However, despite many calls for its inclusion in species threat status (such as the International Union for Conservation of Nature (IUCN) classification system), this has yet to be implemented (Jeon et al. 2024; McLaughlin et al. 2025).

Many European bird populations are in decline (Keller et al. 2020), especially in farmland habitats (Burns et al. 2021), and rare and specialist species such as ground-nesting birds are at increased risk (McMahon et al. 2024; Rigal et al. 2023). The Eurasian curlew (*Numenius arquata*) is a large migratory bird comprising three subspecies, *N. a. arquata*, *N. a. orientalis*, and *N. a. suschkini*, which evolved in refugia during the last glacial period (Tan et al. 2019). The nominate subspecies *N. a. arquata* (hereafter curlew) breeds from Ireland to Spain, the Ural Mountains and up to Arctic Fennoscandia and Russia. The subspecies winters on the Atlantic coast of Western Europe to the North Sea, and south to coastal West Africa (van Gils et al. 2020). The curlew is declining and is listed as Near Threatened by the IUCN (BirdLife International 2021). Population declines in excess of 90% in 40 years have occurred in Irish breeding populations (Balmer et al. 2013; O’Donoghue et al. 2019) in Britain, there has been a 50% reduction in breeding birds between 1995 and 2022 (Heywood et al. 2023); and in Sweden, breeding populations are also declining (Ottosson et al. 2025), with 84% declines between 1976 and 2001 (Wretenberg et al. 2006), and ongoing annual declines of 1.7–1.9% (Green et al. 2026).

Adult survival is high in curlews (Cook et al. 2021; Pakanen and Kylmänen 2023), and declines are instead driven by reduced breeding productivity, from loss of breeding habitat and nest/chick mortality during grassland management (Obłoza et al. 2025; Zielonka et al. 2019). Predation accounts for a high rate of nest failures, 86% in some populations (Ewing et al. 2023; Grant et al. 1999; Zielonka et al. 2019), and can also be the primary cause of fledgling mortalities. To counter this, curlews are the subject of considerable conservation effort across Europe, including habitat management (Colhoun et al. 2022; Ewing et al. 2025). Headstarting is the artificial incubation of eggs with subsequent juvenile release. This removes the high-risk period when eggs and chicks are most vulnerable to predation and destruction from agricultural activities.

Curlews have a wide breeding distribution covering different environmental conditions, which may lead to adaptive divergence between populations. These can include divergence for thermal adaptation, divergence at immune loci to different pathogen and parasite communities, and differences in migration timing. Curlews exhibit behavioural differences across latitudinal gradients in nesting habitat (Bocher et al. 2024). Irish birds are resident, while British birds are resident or winter in Ireland, whereas Swedish breeders migrate to Britain and Ireland for winter (Brown 2015), imposing a higher migratory burden. Weather appears to play little role in curlew migration timing, suggesting a genetic driver (Schwemmer et al. 2021). Birds breeding at higher latitudes consistently depart wintering grounds later, likely timing their arrival with snowmelt (Pederson et al. 2022; Schwemmer et al. 2021).

To date, curlew genetics research has been limited to small segments of mitochondrial and nuclear DNA from curlews in Spain, Germany, Sweden and Russia, revealing that the Iberian (Spanish) population is isolated with low genetic variation, but remains undifferentiated from other populations (Rodrigues et al. 2019). Demographic history analyses have shown a large decline in curlew population size across its range during the Late Quaternary, but more recent losses in genetic diversity were not yet apparent (Tan et al. 2023). However, these studies either included a small number of markers or lacked a strict focus on conservation. Little is known about finer-scale differentiation, inbreeding, and adaptive divergence across curlew breeding habitats. However, a new chromosome-level reference genome (Walsh et al. 2025), now facilitates high-resolution studies using whole-genome resequencing data, allowing for a more complete assessment of the curlew’s current genomic status. Since future conservation actions may include translocating birds between breeding populations, understanding interpopulation genetic diversity and structure is essential.

Here, we evaluated the conservation genetics of curlews using whole-genome single nucleotide polymorphism (SNP) data from 56 individuals across three breeding populations in Ireland, Britain and northern Sweden. We quantifed the four EBVs for all three populations and established baseline data to inform ongoing conservation efforts. We also investigated patterns of genomic differentiation at candidate loci that may reflect either divergent selection or drift among breeding locations.

## Methods

### Study areas

We sampled curlews from Ireland, Britain, and northern Sweden (Fig. 1). In Britain and Ireland, curlews were once common in farmland habitat (Balmer et al. 2013; Sharrock 1976). Today, Irish curlews breed predominantly in peatland (∼74.2%), with smaller numbers in grassland (O’Donoghue et al. 2019). British curlews breed in similar habitats, with strongholds in upland areas (Franks et al. 2017). The Swedish samples were collected from farmland near Umeå in Västerbotten County, from a relatively small area (<200 km^2^). Swedish birds sampled are likely part of a larger, continuously distributed breeding population in both farmland and peatland habitats. Irish samples were collected from across the island, and British samples were collected throughout England, with three samples from Wales.

**Fig. 1.**
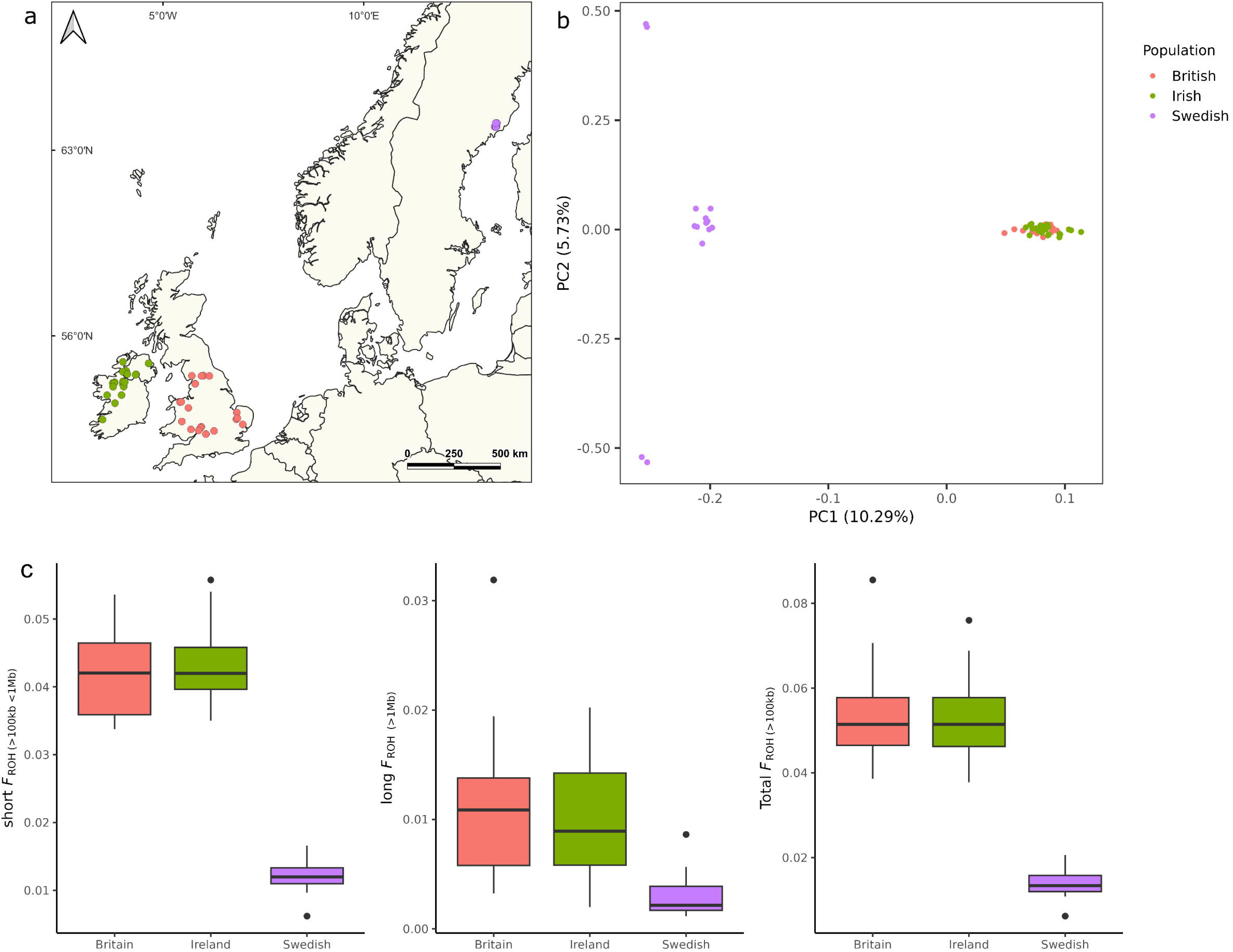
Details of samples from 56 Eurasian curlews (*Numenius arquata*) across three study groups in Ireland, Britain, and northern Sweden showing (a) sampling locations of Eurasian curlew (*Numenius arquata*), with colours indicating the three study populations, (b) the results of the principal component analysis (PCA) based on 5,523,293 SNPs. The PCA plot for the first two principal components shows each individual, colour coded by country of origin. A scree plot of variance explained by the remaining principal component is shown in Fig. S9c, and (c) the proportion of the genome within runs of homozygosity (FROH) for short (>100 kb < 1 Mb), long (> 1 Mb) and total ROH.

### Curlew DNA samples

A total of 66 samples were collected for the current study (Ireland = 26, Britain = 22, Sweden = 18; Fig. 1a.). Where chicks or eggs were sampled, we ensured each sample originated from different clutches. The 26 Irish samples included ∼0.5 ml of blood drawn from the brachial vein of headstarted chicks (*n* = 8). These were stored in EDTA vacutainers at −20 °C. Body contour feathers (*n* = 11) were taken from headstarted chicks. Additional muscle tissue samples (*n* = 7) were collected from deceased Irish birds. There were 18 Swedish hatched eggshell samples collected in the Umeå area of northern Sweden, on the Gulf of Bothnia. British samples were sourced from an existing project at the University of Sheffield, UK and consisted of tissue from 18 embryos and eggshell membranes from 4 eggs.

### Whole-genome resequencing

DNA was extracted using a Zymo Quick-DNA Miniprep Plus kit (Zymo Research Europe, Freiburg, Germany) following the manufacturer’s instructions for the specific tissue types. Library preparation and sequencing were carried out at the National Genomics Infrastructure (NGI) SciLife Laboratory, in Stockholm, Sweden. Additional sequencing libraries for 10 samples were prepared and sequenced by GENEWIZ (Leipzig, Germany). All batches were sequenced on the Illumina NovaSeq X Plus platform. We targeted 18× genome sequence depth and used a 2 × 150 bp paired-end (PE) configuration for sequencing. For additional details on sequencing and sample preparation, see Table S1.

### Quality control and mapping

Raw reads were initially evaluated with FastQC (v0.11.9) (Andrews 2010) to assess initial quality and define trimming parameters. After this, Trimmomatic (v0.39) (Bolger et al. 2014) was used to remove reads with a Phred score below 30, shorter than 50 bp, or containing adapters. Each paired-end read was then mapped to the curlew reference genome (GenBank assembly: GCA_964106895.1) using the Burrows-Wheeler Aligner (BWA) (v0.7.17) (Li 2013), converted to BAM files, and duplicates were flagged with the MarkDuplicates command in Picard v2.27.5 (Broad Institute 2019). Qualimap (Okonechnikov et al. 2016) was used to assess coverage and mapping quality, and samples with low mapping rates (<50%) were removed. Samples with less than 96% mapping rate were cleaned using Kraken 2 (v2.0.7) (Wood et al. 2019) by classifying against the k2_standard_8 database, which includes RefSeq indices for archaea, bacteria, viruses, plasmid genomes, the UniVec database, and the human reference genome. Unclassified reads were retained and remapped.

Variant calling was carried out according to Genome Analysis Toolkit (GATK) (v4.3.0.0) best practices (Van Der Auwera et al. 2013). Briefly, HaplotypeCaller was run with the *-ERC GVCF* option, and individual GVCFs were merged with GenomicsDBImport. Variants were called with the GenotypeGVCFs tool. For each sample, a GVCF was created containing only variant sites and another with all sites. Next, SelectVariants was used to isolate biallelic SNPs and indels. The SNP dataset was filtered using VariantFiltration with the filtering criteria: [QUAL < 30, MQ < 40.00, SOR > 4.000, QD < 2.00, FS > 60.000, MQRankSum < -12.5, ReadPosRankSum < -8.000], and SNPs within 5 bp of an indel were removed with BEDTools (v2.27.1) (Quinlan and Hall 2010).

Only the autosomes were retained, and hard filters were applied using VCFtools, removing any site with a minor allele count (MAC) < 3 or with >10% missing genotypes. A MAC filter was considered more appropriate than the more commonly used minor allele frequency (MAF) threshold, as sample sizes were unequal (Hemstrom et al. 2024). Sites with relatively high depth (>40) or relatively low depth (<10) were also removed. The variant file with only biallelic SNPs was pruned for linkage disequilibrium (LD) at *r*^2^ = 0.2 using PLINK (v1.9) (Purcell et al. 2007) with a 50 SNP window size and 10 SNP step size. The final autosomal SNP dataset was phased with SHAPEIT4 (v4.2.0) (Delaneau et al. 2019) and the default recombination rate of 1 cM/Mb.

The Irish breeding curlew population is extremely small (∼250 pairs island-wide), and the Swedish samples were from a small area. To detect any closely related individuals that could bias downstream population genomics analyses, we ran KING v2.3 (Manichaikul et al. 2010), and 1^st^- degree relatives (kinship coefficient > 0.2) were removed.

### Genetic essential biodiversity variables

We calculated nucleotide diversity (π_SNPs_) using VCFtools with 15-kb windows. VCFtools only considers variant sites and therefore may underestimate genetic diversity, so we also used pixy v1.2.7 (Korunes and Samuk 2021) to estimate genome-wide nucleotide diversity (π_gw_) in 15-kb windows for the all-sites VCF. VCFtools was also used to calculate the observed heterozygosity (*H*_o_). We measured Tajima’s *D* and Watterson’s estimator (θ_W_) using 15-kb windows with pixy (Bailey et al. 2025). As pixy is designed to run on an all-sites VCF, the SNP data set with no minor allele frequency filtering was used to calculate π_gw_, θ_W_ and *D.* We tested whether the mean Tajima’s *D* between populations differed significantly using a block jackknife approach across non-overlapping 1 Mb blocks. Tajima’s *D* was averaged over each block, and pairwise differences between populations were tested using a *t*- statistic derived from the jackknife standard errors.

The *F*_ST_ and *D*_xy_ measures of genetic differentiation were calculated between the populations. For this, an additional comparison was performed using the merged Irish and British populations. The *F*_ST_ is the difference between two sub-populations relative to the total variation within the sampled metapopulation, while *D*_xy_ is the average number of nucleotide differences per site between two populations. Both metrics were calculated using pixy with 15-kb windows.

Inbreeding was estimated using the genomic inbreeding coefficient (*F*) calculated with VCFtools. *F* is excess or deficit of homozygosity within an individual relative to Hardy-Weinberg equilibrium expectations. We also catalogued runs of homozygosity (ROH) with BCFtools/RoH (v1.15.1) (Narasimhan et al. 2016) using a recombination rate of *--rec-rate* = 2 × 10^−8^ (Mugal et al. 2013; Topaloudis et al. 2025). ROHs were identified as homozygous regions longer than 100 kb. From this, *F*_ROH_, the inbreeding coefficient from the ROH detected in each individual curlew (McQuillan et al. 2008) was calculated. The total length of all ROH was calculated, as well as short (>100 kb <1 Mb), and long (>1Mb) ROH. To calculate *F*_ROH,_ the sum of the lengths of ROH in each class was divided by the total length of the autosomal genome for each individual and averaged across populations. Long ROH in particular are informative regarding recent inbreeding, as recombination breaks down ROH over time (Ceballos et al. 2018).

Contemporary *N*_e_ was calculated using ONeSAMP v3.0 (Hong et al. 2024) which is computationally demanding. Therefore, the SNP dataset was thinned for each population using VCFtools with *--thin 1000000* and *--max-missing 1* for the 10 largest chromosomes. This resulted in a dataset consisting of 816 SNPs for Ireland, 818 SNPs for Britain and 819 SNPs for Sweden. We used a mutation rate (μ) of 8.11 × 10^−8^, estimated for *Charadrius* shorebirds (Wang et al. 2019) from the same order (Charadriiformes) as the curlew, which has previously been applied to *Numenius* species to estimate *N*_e_ (Tan et al. 2023).

### Population structure

Principal component analysis (PCA) was conducted using PLINK with the *--pca* option on the dataset of 5,523,293 SNPs, and visualised in ggplot2 v3.3.3 (Wickham 2016) using R. Ancestry and genetic structure were also investigated using ADMIXTURE (v1.3.0) (Alexander et al. 2009) with the LD pruned SNP dataset (*r*^2^ = 0.2). This was run for *K* = 1–5, and the cross-validation error for each *K* value was checked to identify the best-supported number of putative breeding population clusters. Results were visualised with tidyverse (v2.0.0) (Wickham et al. 2019 ) and ggplot2 in R.

### Local adaptation and outlier detection

#### Outlier detection based on composite selection signals

We used Composite Selection Signals (CSS) to investigate natural selection and environmental adaptation using the phased data. This followed the methods as described in Randhawa et al. (2014). Briefly, three summary statistics were calculated between the Irish/British and Swedish samples. These were SNP *F*_ST_ (Weir and Cockerham 1984), *XP*-*EHH* (cross-population extended haplotype homozygosity) and the directional change in the selected allele frequency (Δ*SAF*). These were combined into single per-site CSS scores. These were averaged over 20 kb windows, and SNPs in the top 0.1% of scores that were flanked (± 0.5 Mb) by 5 or more SNPs in the top 1% were considered outliers. The span between flanking SNPs was considered a region of differentiation, and regions within 250kb were merged. Genes within these regions and ± 200 kb around them were extracted from the genome annotation.

#### Outlier detection based on principal components

We also used the R-package pcadapt (v4.4.1) (Privé et al. 2020) to detect divergent selection. This detects outlier SNPs using principal components (PCs) that capture population structure. The first 10 PCs were inspected with a scree plot (Fig S9c), and two PCs relating to population structure were retained. SNPs were regressed against the two PCs, and *z*-scores were calculated; the Mahalanobis distances of which were converted into *P-*values and then transformed into *q*-values (Dabney et al. 2004). Outlier genes were extracted using a false discovery rate (FDR) of 0.01 with the Storey *q*-value method as implemented in the R package qvalue (v2.34.0) (Storey et al. 2023). This analysis was only conducted for the Irish/British vs Swedish comparison as analysis of population structure indicated these were two main clusters. Outlier SNPs were intersected with the genome annotation, and genes ±5 kb of these SNPs were extracted. The Irish and British samples clustered together on the PCA, so this comparison was omitted.

For the functional analysis, both outlier lists were pooled, and gene ontology (GO) term overrepresentation analysis was performed using g:Profiler (Kolberg et al. 2023). Chicken (*Gallus gallus*) was the input query species, as it is the avian species with the most detailed and high-quality functional biological curation and the default g:SCS-adjusted *P*-value (*P*adj) significance threshold of 0.05. Such curations are typically only available for model species. To note, curlew and domestic chicken diverged approximately 91 Myr ago (Kumar et al. 2017), and therefore some curlew-specific biology is likely missed. Nonetheless, this approach is widely used with similar divergence times, for example, for the tree swallow (*Tachycineta bicolor*) (Woodruff et al. 2025) and cattle (*Bos taurus*) (O’Grady et al. 2025), which both used GO terms derived from humans with divergence times of ∼319 Myr and ∼94 Myr respectively (Kumar et al. 2017).

## Results

### Sequencing, quality control and relatedness

Low mapping rate (MR) values required three samples to be removed from the SNP/indel genotyping process and downstream population genomics analyses: one Irish (MR = 35%), one British (MR = 13%), and one Swedish (MR = 54%). Many of the samples originated from embryo tissues sampled from the wild or from similarly deteriorated tissue. This resulted in 11 samples having low mapping rates, ranging from 59% to 96%. By filtering contamination with Kraken 2, the mapping rates increased to 97–98% for these samples, except for one British sample (82%), which was removed. An additional British sample had a spurious GC-content and was also removed, with a final total of 61 curlew samples retained for SNP/indel genotyping.

Heatmaps showing kinship within each study population (Fig. S1–3) were produced to identify any highly related individuals. This revealed four sets of 1^st^-degree relatives (kinship coefficient > 0.2), two in Ireland and two in Sweden. The four with the lowest coverage were removed. This analysis also revealed one British sample was highly divergent and it was removed. These samples were then joint genotyped for SNPs/indels again resulting in the final dataset consisted of 56 samples (Ireland = 23, Britain = 18, Sweden = 15). Of the final Irish samples, 8 originated from blood, 6 were from muscle tissue, and 9 from feathers. The British samples consisted of 15 embryonic tissues and 3 eggshell membranes.

### Genome-wide SNP numbers

Table S2 shows the number of genome-wide SNPs obtained for the different datasets. The final total for the main dataset was 5,523,293 SNPs, and after LD-pruning, 782,547 SNPs were retained. For the indels, 2,257,915 passed the filtering criteria. The mean individual sample sequencing depths (±SD) for each population post-filtering were: Ireland (*n* = 23): 17.752 ± 3.954×; Britain (*n* = 18): 19.179 ± 3.734×; and Sweden (*n* = 15): 17.152 ± 1.514×. The mean sequencing depth across all 56 samples was 18.050 ± 3.438×.

### Genetic diversity

Genome-wide nucleotide diversity was similar across all three populations (Table 1). Watterson’s θ_W_ and observed heterozygosity (*H*_o_) are also similar across the three populations, but Tajima’s *D*, calculated from SNPs and monomorphic sites, differed somewhat. Tajima’s *D* was positive in the Irish and British populations (Ireland: 0.200 ± 0.007; Britain: 0.136 ± 0.006, block jackknife SE) but negative in Sweden (-0.216 ± 0.006, block jackknife SE), and all pairwise differences were statistically significant (block jackknife t-test, all *P*-values < 0.001).

**Table 1.** Mean population-level indices of genetic variation (+ SD) and inbreeding estimates for Eurasian curlew (*Numenius arquata*) across three study populations (Ireland, Britain, and Sweden).

|  | I | r | e | I | a | n | d | B | r | i | t | a | i | n | S | w | e | d | e | n |
| --- | --- | --- | --- | --- | --- | --- | --- | --- | --- | --- | --- | --- | --- | --- | --- | --- | --- | --- | --- | --- |
| $H_o$ | 0.256 | | | | | | ± 0.003 | 0.255 | | | | | | ± 0.002 | 0.258 | | | | | ± 0.002 |
| $\pi_{\text{SNPs}}$ | 0.001 | | | | | | ± 0.001 | 0.001 | | | | | | ± 0.001 | 0.001 | | | | | ± 0.001 |
| $\pi_{\text{gw}}$ | 0.003 | | | | | | ± 0.002 | 0.003 | | | | | | ± 0.002 | 0.003 | | | | | ± 0.002 |
| $\theta_w$ | 0.002 | | | | | | ± 0.002 | 0.002 | | | | | | ± 0.002 | 0.003 | | | | | ± 0.002 |
| Tajima's $D$ (SE) | 0.200 | | | | | | ± 0.007 | 0.136 | | | | | | ± 0.006 | -0.216 | | | | | ± 0.006 |
| $N_e$ (95% CI) | 193 | | | | | | ± 148-244 | 186 | | | | | | ± 145-236 | 189 | | | | | ± 145-236 |
| $F_{\text{ROH}}$ | 0.058 | | | | | | ± 0.011 | 0.059 | | | | | | ± 0.009 | 0.018 | | | | | ± 0.004 |
| Long $F_{\text{ROH}}$ | 0.012 | | | | | | ± 0.006 | 0.012 | | | | | | ± 0.006 | 0.003 | | | | | ± 0.002 |
| $F$ | 0.012 | | | | | | ± 0.012 | 0.016 | | | | | | ± 0.009 | 0.003 | | | | | ± 0.006 |

Contemporary effective population size (*N*_e_) does not differ significantly (95% confidence intervals overlap). Ireland showed the highest contemporary *N*_e_ (193, CI: 148–244), followed by Sweden (189, CI: 145–236) and Britain (186, CI: 145–236).

Both measures of inbreeding (*F* and *F*_ROH_) were higher in the Irish and British breeding curlew samples than in the Swedish samples (Table 1; Fig. 1c). For each ROH class (small, long, and total), the Irish and British breeding populations show higher levels of inbreeding than the Swedish samples. The estimates of *F*_ROH_ from long ROH (>1 Mb) were identical for the Irish and British populations: *F*_ROH_ (± SD) = 0.012 ± 0.006 in both populations. For the Swedish population, this measure of inbreeding was approximately fourfold lower: *F*_ROH_ = 0.003 ± 0.002. The *F*_ROH_ for short and long ROH showed the same trend. This difference is unlikely to reflect sampling scale, as the confined Swedish sampling would tend to inflate rather than depress Swedish *F*_ROH_.

### Population structure

Interpopulation genetic diversity was evaluated among all three breeding populations using *F*_ST_ values, and for a combined Ireland-and-Britain group versus Sweden (Table 2). A very small *F*_ST_ value of 0.0018 was observed between the Irish and British curlew populations, compared with pairwise comparisons for the Swedish population: Ireland versus Sweden, 0.032; Britain versus Sweden, 0.031; and Ireland and Britain versus Sweden, 0.032. The alternative measure of interpopulation genetic diversity (*D*_xy_) was consistent across all population comparisons (∼0.003) (see Table 2).

**Table 2.** Pairwise genetic differentiation (*F*_ST_) and absolute divergence (*D*_xy_) among Eurasian curlew (*Numenius arquata*) populations.

| $F_{ST}$ | Ireland | Sweden |
| --- | --- | --- |
| Ireland |  | 0.032 |
| Britain | 0.002 | 0.032 |
| Ireland-Britain |  | 0.032 |
| $D_{xy}$ | | |
| Ireland |  | 0.003 |
| Britain | 0.003 | 0.003 |
| Ireland-Britain |  | 0.003 |

The analysis of genetic structure and ancestry performed with ADMIXTURE (Fig. 2) is consistent with the PCA result. For *K = 2*, which showed the lowest cross-validation error (0.53) except for *K* = 1 (Fig 2b), two distinct clusters are evident, one for the Irish and British birds and one for the Swedish birds. At *K* = 3, genetic structure within the Ireland and Britain group is detected, and for *K* = 4, additional structure in the Swedish population emerges. However, PC2 shows the Swedish population separating more than the Irish and British cohort.

**Fig. 2.**
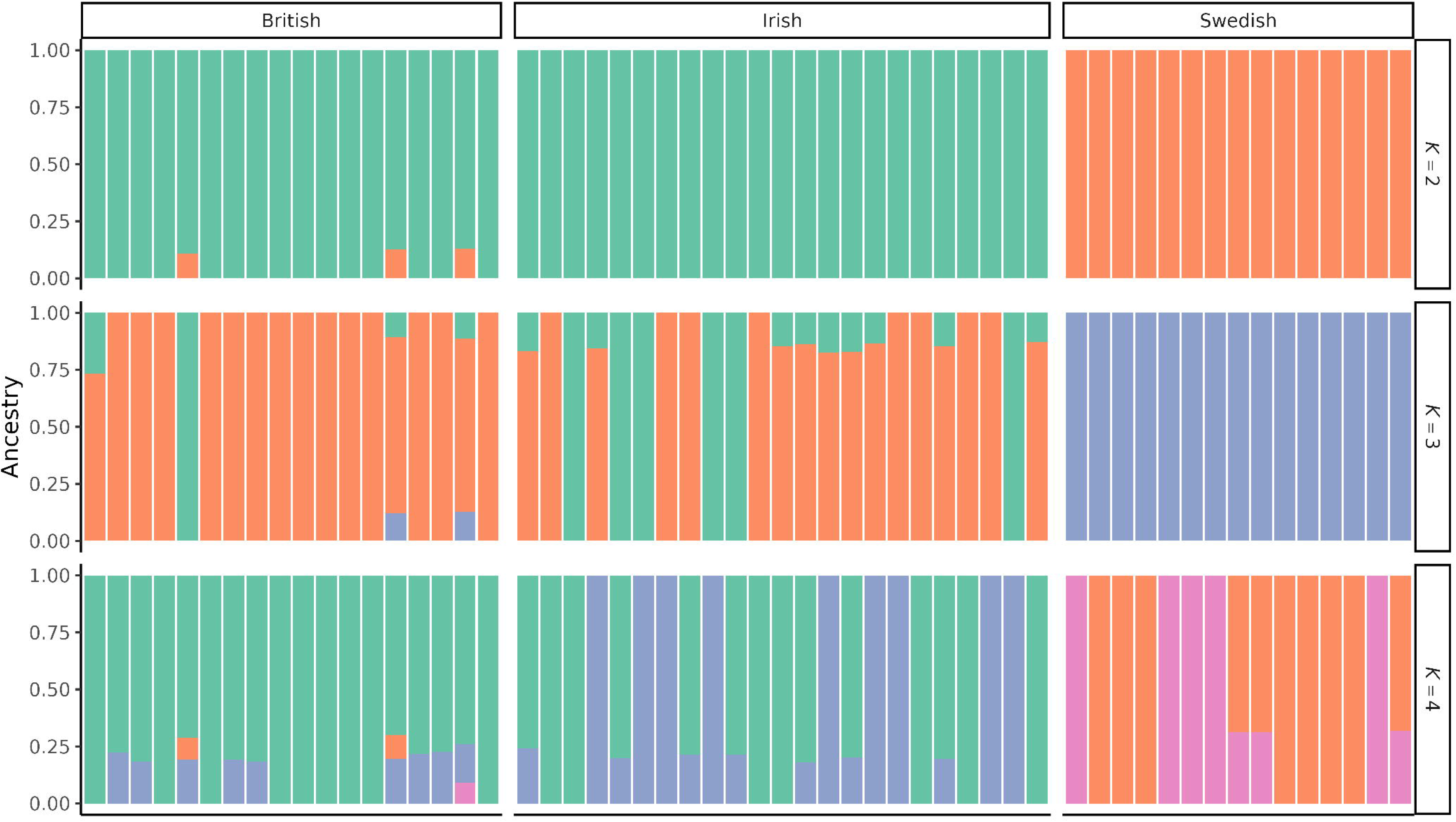
Plot of ADMIXTURE results for 56 Eurasian curlews (*Numenius arquata*) across three study groups in Ireland, Britain, and northern Sweden. based on 782,547 SNPs. The three plots show estimated individual ancestries for each individual sampled for *K* = 2 to *K* = 4 ancestral populations. The cross-validation (CV) error values show that *K* = 2 has the lowest CV-error, except for *K* = 1 (Fig. S10).

### Selection signals

#### Composite selection signals

The CSS analysis comparing the combined Irish and British curlew group and the Swedish group identified 34 regions of putative differential selection (Fig. 3). Of these regions, 30 contained 198 annotated genes. The CSS regions were distributed across 13 chromosomes, with chromosome 2 (NCBI RefSeq: NC_133577.1) containing the most regions (6). The list of outlier genes and outlier regions is detailed in Appendix 2. A CSS analysis was also performed for the Irish and British curlew populations (Fig. S7), with 37 regions identified containing 369 genes. This is somewhat counterintuitive and likely due to high noise from the overall low level of differentiation.

**Fig. 3.**
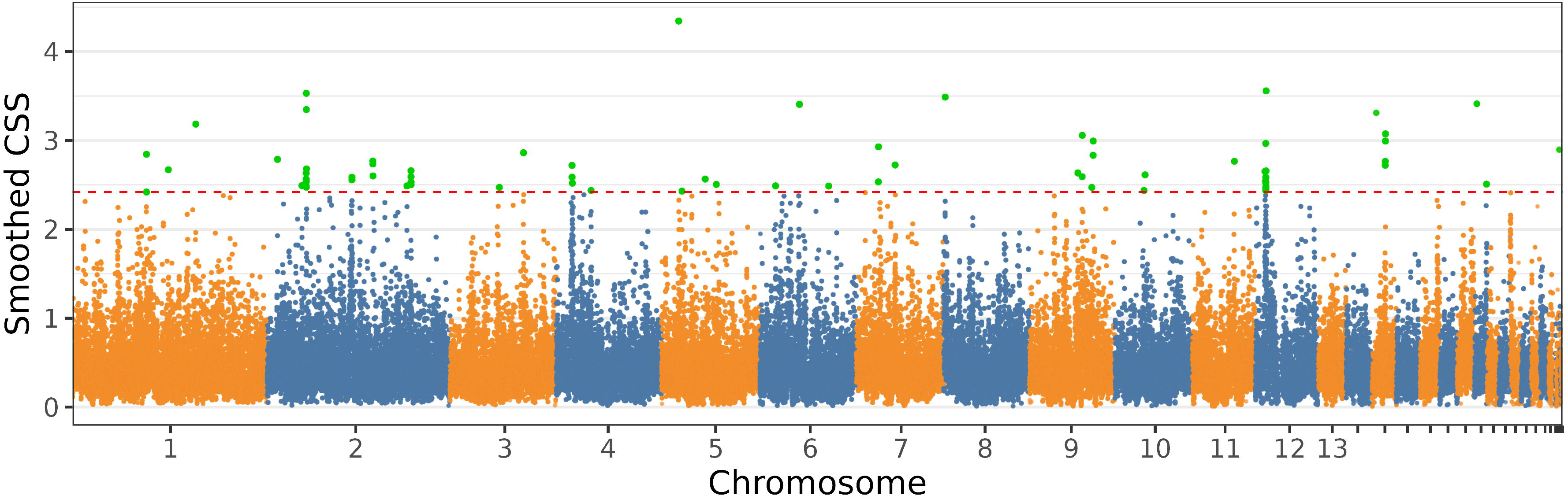
Composite Selection Signal (CSS) scores across the genome for Irish and British versus Swedish Eurasian curlew (*Numenius arquata*) populations. Peaks indicate regions under putative divergent selection based on combined metrics (*F*_ST_, *XP-EHH*, and *ΔSAF*) smoothed in 20 kb windows. The dashed red line represents the genome-wide 0.1% threshold of the CSS scores. The green points are SNPs above the 0.1% threshold and also flanked by at least 5 SNPs in the top 1% of CSS scores.

#### Principal component selection signals

Using pcadapt, we identified 24,692 outlier SNPs. From the reference genome annotation, we extracted 612 genes that either contained outlier SNPs or were located ±5 kb of a gene body. Of these, 388 were unique, and 270 had assigned gene symbols available. Outlier SNPs were identified within 162/388 unique genes. For the remaining 226 genes, SNPs were located 5 kb upstream or downstream and could, therefore, be located in gene regulatory elements (GREs). This is likely a result of linkage disequilibrium effects with multiple SNPs mapping to the same gene. The full outlier gene list is shown in Appendix 3.

### Outlier genes and gene ontology (GO) overrepresentation analysis

After removing duplicates between pcadapt and the CSS methods, there were 432 unique gene names. The resulting GO term plot, generated by g:Profiler, is shown in Fig. S5; for further GO term overrepresentation results, see Table S3. There were five significant GO terms, three in the Molecular Function category (*G protein-coupled peptide receptor activity*, *peptide receptor activity* and *peptide binding*) and two in the Cellular Component category (*cytosolic ribosome*, *ribosomal subunit*). Three genes overlapped between the two outlier detection methods. These were *MC3R* (melanocortin 3 receptor), *FLRT3* (fibronectin leucine rich transmembrane protein 3) and *NPBWR1* (neuropeptides B and W receptor 1).

## Discussion

Here, we provide the first exploratory population genomics study of three breeding localities of Eurasian curlew in Ireland, Britain, and Sweden, reporting the Essential Biodiversity Variables for these populations and assessing functional divergence among them. Using whole-genome data from 56 curlews, we show early genomic signals of population declines, including increased inbreeding and a slightly elevated Tajima’s *D* in Irish and British breeding curlew.

### Genetic diversity in the Irish, British, and Swedish breeding Eurasian curlew populations

The rate of population decline across the three study populations varies (Green et al. 2026; Heywood et al. 2023; O’Donoghue et al. 2019; Ottosson et al. 2025); however, genetic diversity across the three populations was similar for many metrics (π, *H*o, and contemporary *N*_e_). These population declines are recent (within the last 50 years), and it can take some time for changes in genetic diversity to be detected via genomic approaches. Genome-wide nucleotide diversity (π) is a lagging measure of genetic diversity (Jeon et al. 2024), impacted by ancient demography rather than ongoing bottlenecks. Watterson’s Theta (θ_w_), a leading indicator, is impacted more by contemporary *N*_e_ (Jeon et al. 2024; Tajima 1989a) as it is sensitive to the loss of rare alleles, and excess of intermediate-frequency variants due to a bottleneck (Gattepaille et al. 2013). Tajima’s *D* is calculated from π_gw_and θ_w_ (Tajima 1989b). A negative *D* value indicates population growth; if *D* ≈ 0, the population is stable, and a positive *D* value can indicate population decline or a recent bottleneck due to an excess of intermediate-frequency variants. Estimates of *D* in this study were positive for the Irish and British samples, but negative in Sweden. The higher *D* value observed in the Irish curlew samples likely reflects a more severe breeding population contraction (O’Donoghue et al. 2019), which is consistent with the threatened status of the Irish breeding curlew. These results are consistent with early-stage genomic effects of population decline: whether diversity could recover depends on future demographic trajectories.

The discrepancy in expected genetic diversity, i.e. uniform across populations, could reflect differences in sampling distribution. The Swedish samples came from a confined area of Sweden, and our results likely underestimate the genetic diversity in Sweden, whereas the Irish and British samples were derived from across the current breeding range and are likely more representative. The time lag that occurs between population decline and the loss of genetic diversity (Gargiulo et al. 2025; Liu et al. 2025) —termed ‘genetic extinction debt’—should also be considered. Curlews are more susceptible to this due to several life history traits, including a long generation interval of 9.5 years (Bird et al. 2020), corresponding to ∼ 4.5 generations since severe declines began in Ireland, long lifespan, overlapping generations, high survival (apparent for adult curlew), and a relatively large historical population size with a wide distribution (Gargiulo et al. 2025). This means that if population sizes were to stabilise, genetic diversity would not necessarily stop decreasing; it would likely continue. However, this time lag can give practitioners time to act (Gargiulo et al. 2025) and implement habitat restoration to improve nesting habitat and reduce the abundance of generalist predators, so it should also be considered when monitoring population recovery and status.

We show the onset of genetic diversity loss due to inbreeding. Inbreeding, measured as long ROH, *F*_ROH >1 Mb_, is higher in the Irish and British populations, relative to the breeding curlews sampled in Sweden. The *F*_ROH >1 Mb_ for the British and Irish populations was approximately 0.012, compared to 0.003 in Sweden. Kyriazis et al. (2025) considered an *F*_ROH_ of 0.02 to represent low, to no inbreeding, in the passerine akikiki (*Oreomystis bairdi*). Low levels of inbreeding, with no ROH > 0.5 Mb despite large-scale population declines were recorded in a recently declined population of regent honeyeater (*Anthochaera phrygia*), potentially attributed to high mobility and historically large populations (Liu et al. 2025) resulting in a lag in genomic effects being detectable. High *F*_ROH_ and inbreeding depression occur in the wild red-headed wood pigeon (*Columba janthina nitens*) with a long *F*_ROH >1 Mb_ of 0.202, following a severe and prolonged bottleneck (Tsujimoto et al. 2025). Therefore, while *F*_ROH_ is not extreme in any of our populations, the elevated levels in Ireland and Britain indicate an increased vulnerability to inbreeding-related effects, should declines continue. In addition, the confined Swedish sampling would be expected to inflate Swedish ROH estimates. We see the opposite pattern (lowest *F*_ROH_ in Sweden) suggesting the elevated Ireland/Britain inbreeding is robust to this confound. In addition, the classical inbreeding coefficient (*F*) shows the same pattern (Table 1).

### Population structure

Irish and British curlew populations are genetically highly similar (*F*_ST_ = 0.002), whereas both populations differ more from the Swedish samples (*F*_ST_ = 0.032). Principal component analysis and analysis of genetic structure and ancestry using ADMIXTURE show the same pattern of population clustering, with some residual admixture from the Swedish breeding population into the British. Given the shorter geographic distance, more gene flow could be expected between Britain and Sweden than between Ireland and Sweden. Swedish birds winter in Britain (Brown 2015), potentially explaining the modest admixture exhibited by three individuals; however, they also winter in Ireland. Another likely driver of population structure for curlews is high natal (Bainbridge and Minton 1978), wintering (Schwemmer et al. 2016; Schwemmer et al. 2021), and breeding (Pakanen and Kylmänen 2023; Valkama et al. 1998) site fidelity. Similar philopatry has led to differentiation in the dunlin (*Calidris alpina*) (Rönkä et al. 2021) and American oystercatcher (*Haematopus palliatus*) (Avila-Cárdenas et al. 2025). With this split, populations can diverge through genetic drift and adaptive selection if environments differ.

### Selection signals and local adaptation

We identified 432 candidate genes under putative divergent selection between Irish/British and Swedish breeding curlew populations using two methods. Given the low *F*_ST_ values among the populations (a maximum of ∼ 0.03), strong adaptive divergence is unlikely. Nonetheless, divergence could be important for conservation, as the outlier genes identified encode proteins involved in ecologically relevant traits in other bird species and vertebrates. These candidate genes should be viewed as working hypotheses that can be tested using additional omics data (e.g., transcriptomics) and/or functional validation in shorebird populations. Here we discuss genes considered important from a conservation viewpoint; for more discussion of outlier genes and traits, see Appendix 1.

Three genes were detected by both the CSS and pcadapt outlier methods. These were *MC3R* (melanocortin 3 receptor), *FLRT3* (fibronectin leucine rich transmembrane protein 3) and *NPBWR1* (neuropeptides B and W receptor 1). The *MC3R* gene encodes a conserved regulator of energy balance and feeding behaviour in birds (Zhang et al. 2017) (McConn et al. 2019), and is linked to circadian entrainment of feeding in mammals (Begriche et al. 2009). Seasonal cycles in food intake occur in birds with regard to migration, and differences in migration distance between the study populations could also be a factor associated with divergence at this locus. Fibronectin-leucine-rich transmembrane (*FLRT*) proteins are associated with embryonic development in mammals (Karaulanov et al. 2006) and chick limb development (Smith and Tickle 2006; Tomás et al. 2011). The *NPBWR1* gene is expressed in the brain and pituitary of chickens; however, its function is unclear (Bu et al. 2016), with a potential role in egg production and ovarian development in adult Wanxi White geese (*Anser cygnoides*) (Li et al. 2024).

Several outlier genes were associated with energetics and appetite. Curlews in Sweden have a shorter breeding season (Pederson et al. 2022) meaning chicks may be under selection for higher energy intake and quicker maturation. Therefore, differential selection could affect genes and GREs associated with energetics, appetite and fattening. The most overrepresented GO term was G protein coupled peptide receptor (GPCR) activity, which is involved in vertebrate feeding and energy homeostasis (Shioda et al. 2008). We also find several outlier genes involved in appetite, including *MC3R* and *CCKAR* (Aderibigbe et al. 2022). Another explanation for some of these genes is selection pressure from a longer migration distance for Swedish curlews along the east-Atlantic flyway, relative to Irish and British birds (Brown 2015). *MC3R* also has a role in circadian timing, as discussed above, as does the outlier gene *ID2* (Ward et al. 2010).

At least 16 immune-related genes were also identified, though no GO term overrepresentation was found. Divergence at immune loci is important for conservation and supplementation to understand the risks of translocating individuals and also potential benefits from introducing novel variation. Immune genes respond fast to divergent selection (Shultz and Sackton 2019) and diversity at these genes can be lost quickly during bottlenecks (Belasen et al. 2019; Cortazar Chinarro et al. 2025). Disease challenges in British waders remain poorly characterised (Beckmann et al. 2025), so it is conceivable that pathogen and parasite burdens differ between the two populations. However, it must be said that these populations are in close proximity at wintering grounds for a large portion of the year and there could be mixing of parasites and pathogens there. Contrary to this, there are pathogens and parasites during the egg and chick stages that may be habitat- and latitude-specific, and selection at these earlier stages could result in divergence. Notably, the *TLR2* gene, which encodes a pathogen-recognition receptor (PRR) (Wigley 2013) was identified as an outlier, and TLR gene allelic diversity can vary among bird populations at small spatial scales (Grueber et al. 2017) and is associated with survival in some endangered bird populations (Bateson et al. 2014; Grueber et al. 2013; Hartmann et al. 2014).

Mean, minimum, maximum, and variability in temperature can all be selective pressures on birds (Nord et al. 2026), with temperature variability being a plausible driver between these study populations. Some notable genes were recorded as outliers. We identified the *CIDEA* gene as an outlier, which has a role in mammalian thermoregulation (Zhang et al. 2014) and is upregulated in chicken in response to cold acclimation (Wang et al. 2025). Four outlier genes identified in this study overlap with genes significantly differentially expressed in tree swallow (*Tachycineta bicolor*), in response to increased temperature (*MITD1*, *TLR2*, *SMARCA1,* and *PCMTD1* ) (Woodruff et al. 2025) and three genes differentially expressed with decreased temperature in domestic turkeys (*Meleagris gallopavo*) (Reed et al. 2025). However, in the absence of thermoregulatory pathway overrepresentation and with many of the genes being stress-, metabolic-, and immune-related in nature, thermal selection remains a potential driver of divergence requiring functional validation.

Several genes with roles in embryogenesis were also identified. This is noteworthy as most of the British samples were from embryos where the cause of failure is unknown. It is also noteworthy that the majority of Irish samples came from headstarted birds, where selection on early life stages is relaxed in the current generation, relative to Swedish birds which are fully wild. Therefore, any apparent divergence at embryo-related loci could be a sample-type artefact, a conservation management artefact, or adaptation to divergent environments. *F*_ST_ outlier distributions did not differ between embryo and non-embryo British and Irish samples, and GO enrichment analysis recovered no terms associated with embryogenesis or viability, suggesting embryo sampling is not driving the developmental genes identified.

## Conclusions

Our study calculated the four genetic EBVs, providing the first genomic baseline of Eurasian curlew breeding in Ireland, Britain and northern Sweden. Despite large-scale declines, especially in Ireland, genetic diversity remains broadly similar across the populations. This may be influenced by the scale and distribution of sampling, where the Swedish sampling distribution is restricted to a small area in northern Sweden, potentially underestimating genetic diversity for the Swedish samples. The marginally elevated Tajima’s *D*, strongest in Ireland, along with higher *F* and *F*_ROH_ values in Ireland and Britain, potentially indicates the early onset of diversity loss and a rising inbreeding risk, raising concerns for these populations. This highlights the genetic lag that may be occurring in these populations between declines and detectable genetic responses. Population structure analyses revealed that Britain and Ireland are near panmictic, but divergent from Swedish breeding populations. Genetic monitoring of curlew populations should be implemented across their range, and suitable material can be collected due to active conservation management in all populations and the wide variety of available samples. The continuous monitoring of these EBVs, ideally with the addition of genomic data from museum specimens, will be required to monitor genomic erosion over time in breeding curlew populations. This will facilitate conservation actions aiming to balance protecting population sizes and long-term evolutionary potential.

Outlier analyses suggest a potential adaptive divergence in energetics, thermal adaptation, and immune function. These findings could have direct implications for translocation and supplementation. Ideally, an investigation into genetic load in Britain and Ireland, along with a larger sample set of viable British samples, would confirm the suitability for translocations. The findings in this study show that conservation-motivated translocations between Britain and Ireland appear appropriate, but in the absence of increased sampling, any transfers involving Swedish curlews to these populations should be approached with caution due to both genetic differentiation and putative local adaptation.

## Acknowledgements

We would like to thank those who provided samples for the study, including Barry O’Donoghue (National Parks and Wildlife Service (NPWS) and Katie Gibb and Amy Burns (both Royal Society for the Protection of Birds). Alyn Walsh (NPWS) assisted with sample collection and provided licensing advice; Jess Hodnett (Fota Wildlife Park) assisted with sample collection; Owen Murphey (BreedingWaders EIP) provided support for the study. We would also like to thank Dr John Browne (UCD) for advice and guidance on sample collection, DNA extractions and sequencing. We are grateful to everyone who contributed to this project. The authors acknowledge support from the National Genomics Infrastructure in Stockholm funded by Science for Life Laboratory, the Knut and Alice Wallenberg Foundation and the Swedish Research Council, and NAISS/Uppsala Multidisciplinary Center for Advanced Computational Science for assistance with massively parallel sequencing and access to the UPPMAX computational infrastructure.

## Author contributions: CRediT

**Grace Walsh:** Conceptualization, Formal analysis, Funding acquisition, Investigation, Methodology, Visualization, Writing – original draft, Writing – review & editing. **Jacob Höglund:** Conceptualization, Funding acquisition, Resources, Supervision, Writing – review & editing. **Patrik Rödin-Mörch:** Conceptualization, Methodology, Software, Writing – review & editing. **James A Ward:** Conceptualization, Methodology, Software, Writing – review & editing. **Rebecca C Örnberg:** Investigation, Writing – review & editing. **Jamie Thompson:** Investigation, Writing – review & editing. **Declan O’Donovan:** Investigation, Methodology, Resources, Writing – review & editing. **Adriaan de Jong:** Conceptualization, Investigation, Resources, Writing – review & editing. **Seán BA Kelly:** Investigation, Methodology, Resources, Writing – review & editing. **Nicola Hemmings:** Resources, Writing – review & editing. **David E MacHugh:** Conceptualization, Data curation, Methodology, Resources, Writing – review & editing. **Barry J. McMahon:** Conceptualization, Funding acquisition, Project administration, Resources, Supervision, Writing – review & editing.

## Data availability statement

The raw sequence data generated for this study is part of an ongoing doctoral project. The thesis will be submitted shortly and all data will be uploaded to the Sequence Read Archive, and will be uploaded in advance of any publication.

## Ethics and licencing statement

Blood samples were taken under licence from the National Parks and Wildlife Service (NPWS) (licence no. C154/2022), approved by the Animal Research Ethics Committee at University College Dublin, Ireland (Approval no. AREC-22-27-McMahon), and project authorisation was received from the Health Products Regulatory Authority (Project authorisation: AE18982/P231). Body contour feathers (*n* = 11) were taken from headstarted chicks under licence from the NPWS (licence no. C104/2023).

