## Appendix 1 for "Genomic status of the Eurasian curlew *Numenius arquata*: estimating Essential Biodiversity Variables and selection signals for a declining migratory bird"

**Appendix 1: Supplementary Methods and Discussion**

Supplementary Tables and Figures

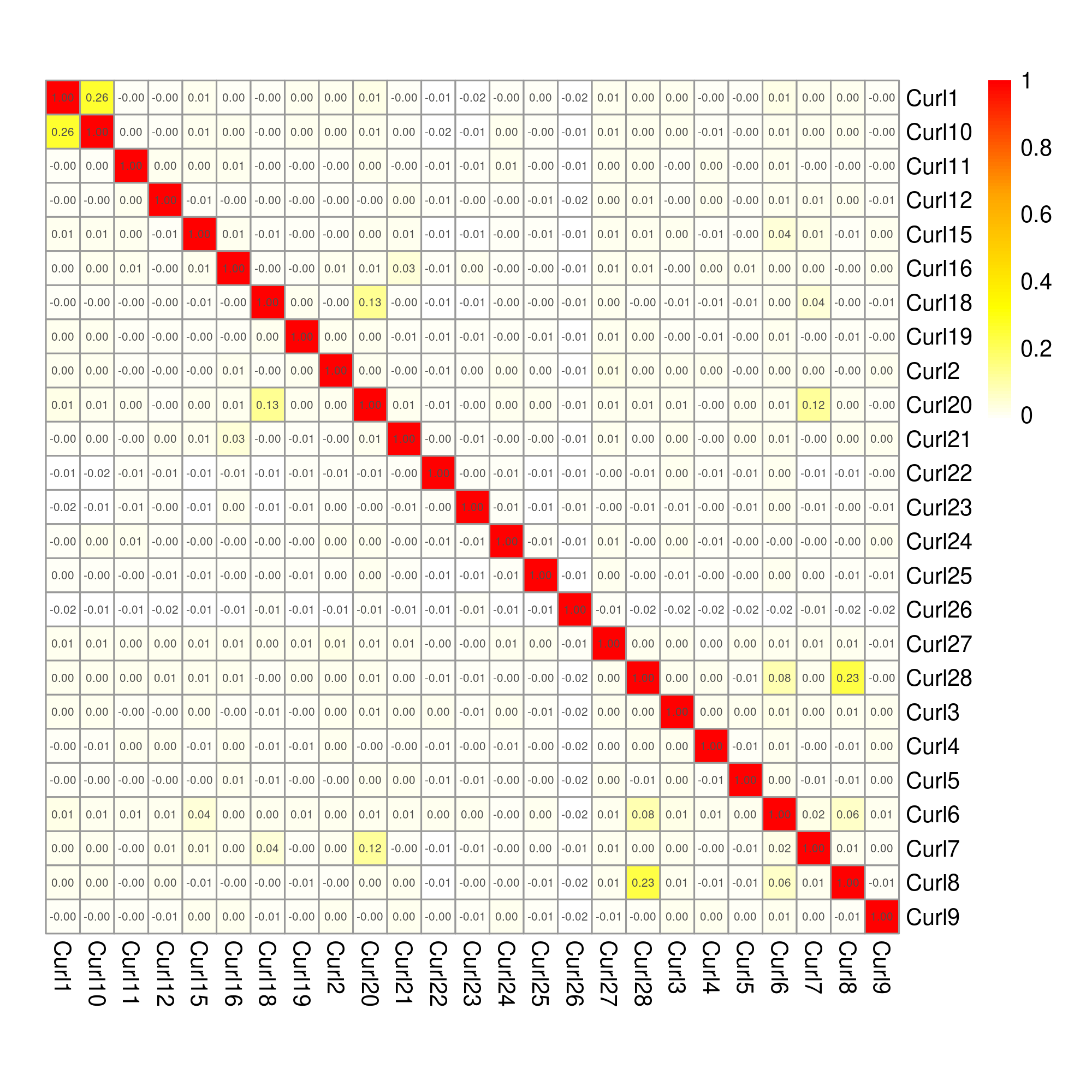

**Fig. S1.** Heatmap showing kinship coefficients for 25 Irish curlews generated with KING 2.2.7.

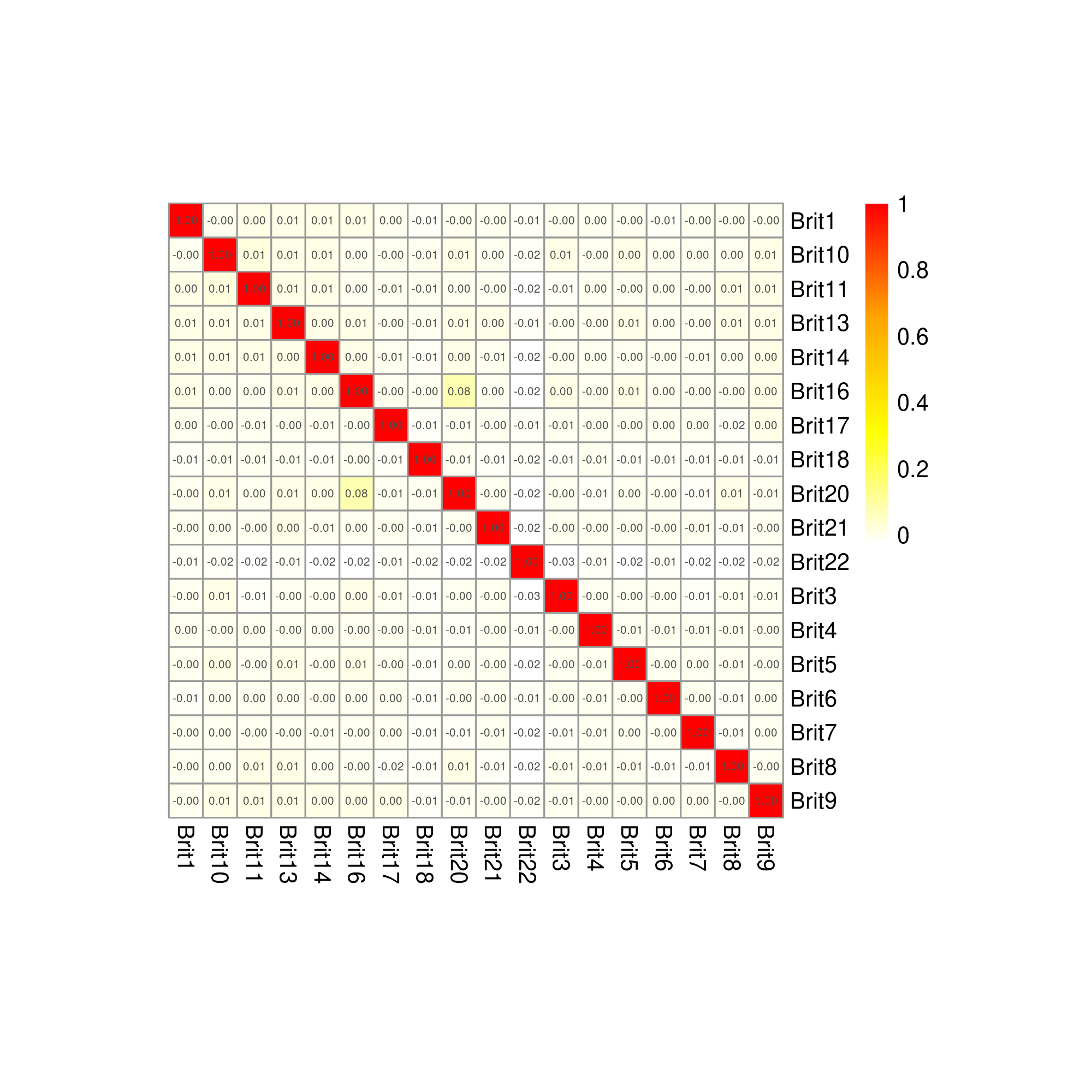

**Fig. S2.** Heatmap showing kinship coefficients for 18 British curlews generated with KING 2.2.7.

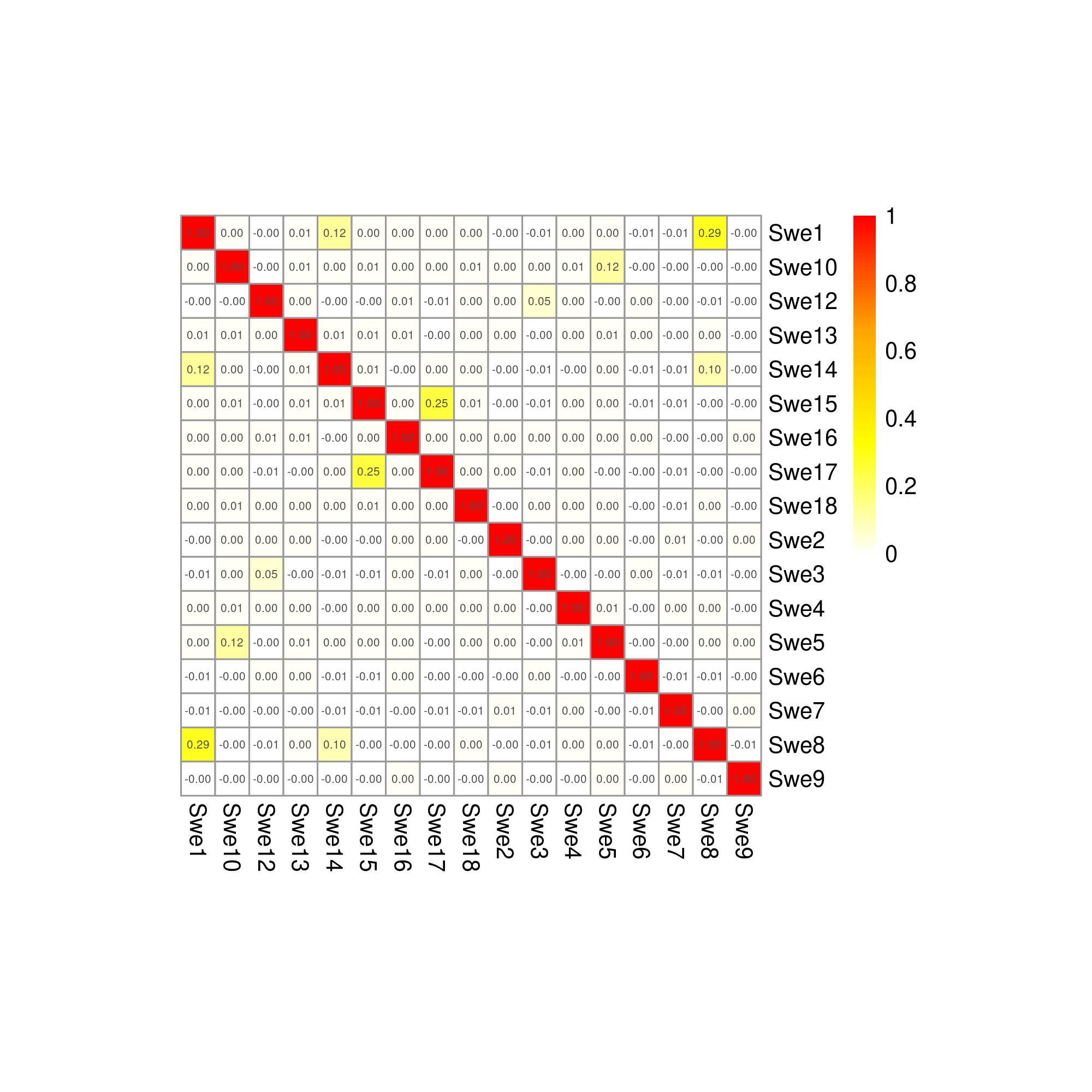

**Fig. S3.** Heatmap showing kinship coefficients for 17 Swedish curlews generated with KING 2.2.7.

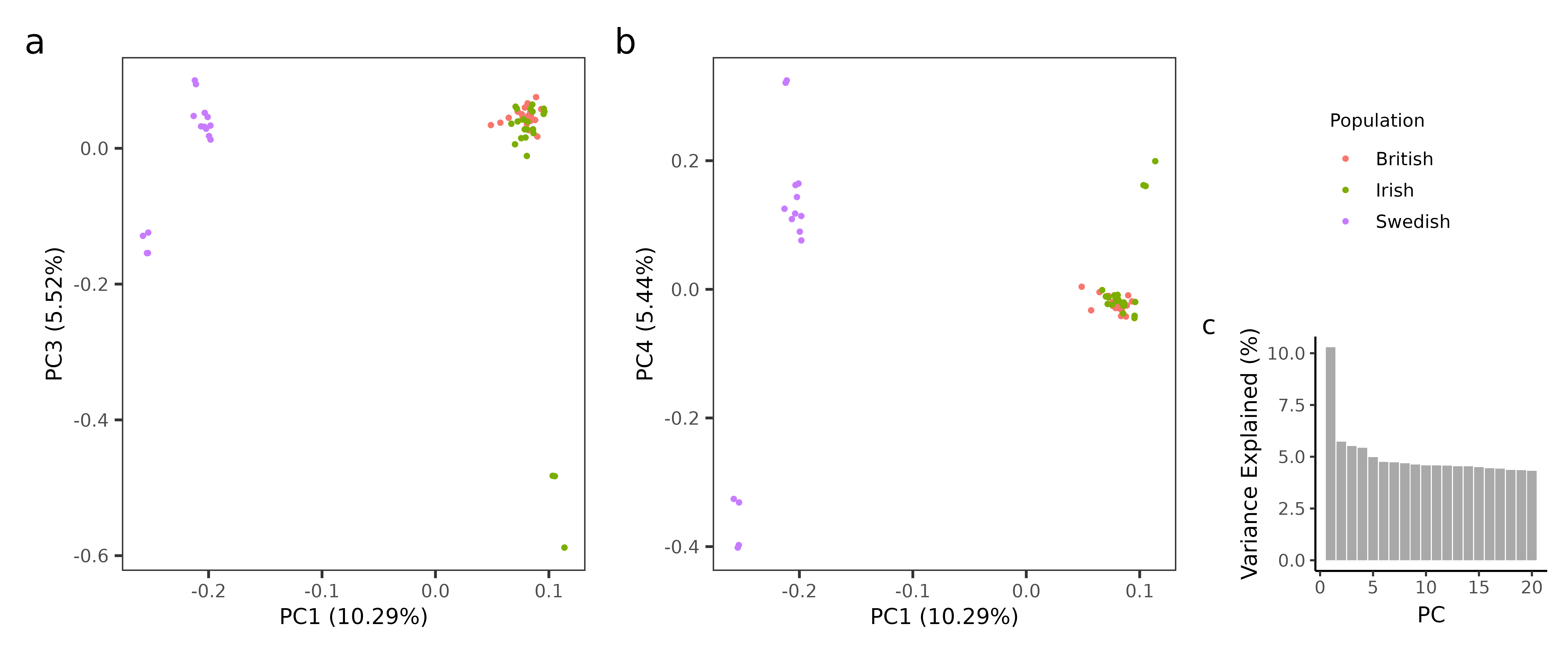

**Fig. S4.** Results of the principal component analysis (PCA) for 56 Eurasian curlews (*Numenius arquata*) across three study groups in Ireland, Britain, and northern Sweden based on 5,523,293 SNPs. The plots show (a) principal component 1 versus 3, (b) principal component 1 versus 4 and (c) a histogram of the variance explained by the first 20 principal components.

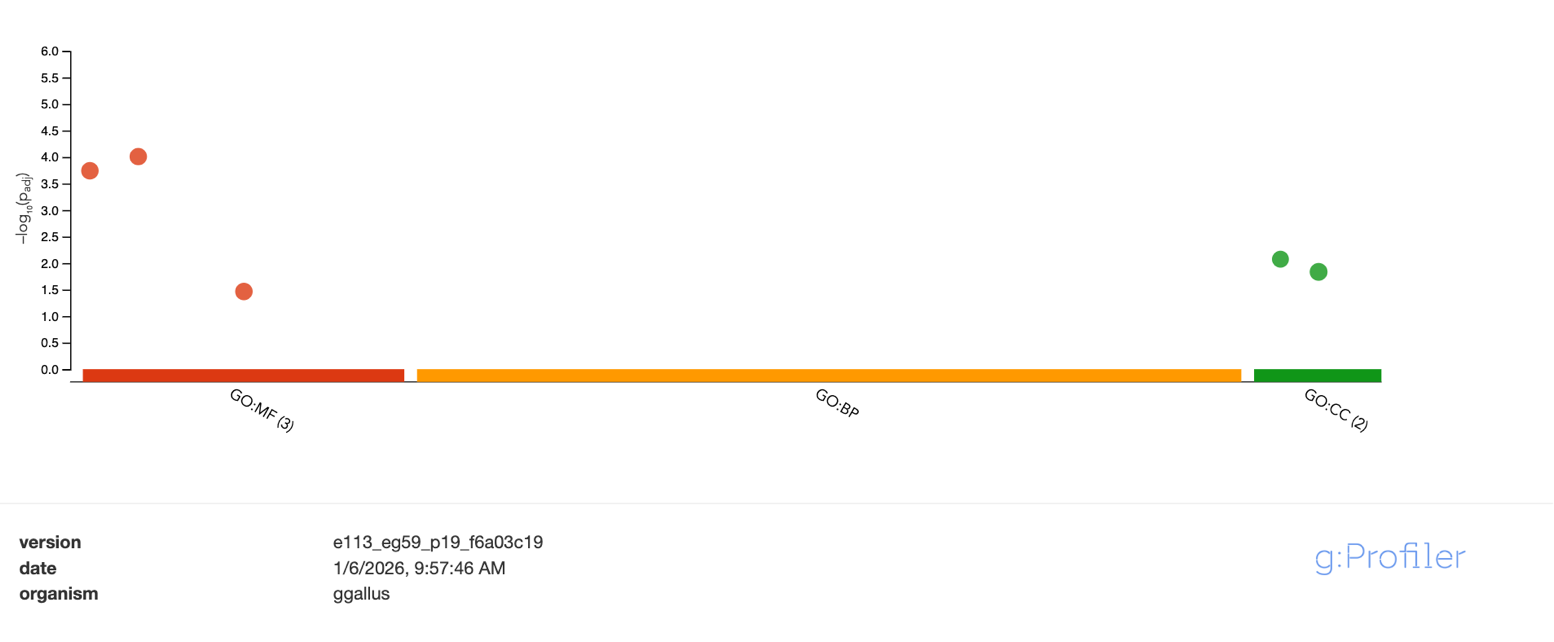

**Fig. S5.** Gene Ontology (GO) overrepresentation analysis for the outlier genes (n=432) identified using the R package pcadapt and Composite Selection Signal (CSS) scores for Irish/British versus Swedish Eurasian curlew (*Numenius arquata*).

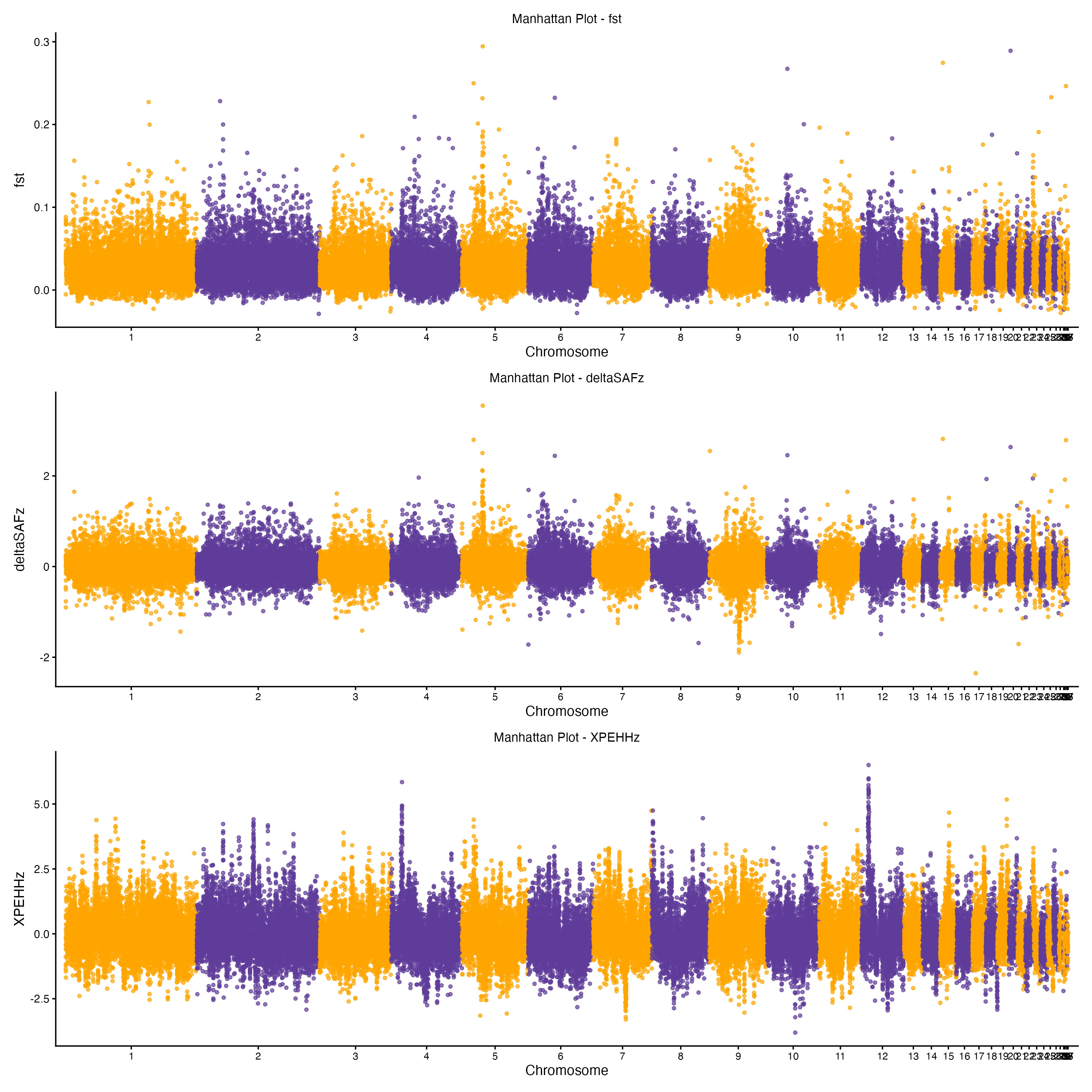

**Fig. S6.** Manhattan plots of three summary statistics (*F*_ST_, *ΔSAF* and *XP-EHH*) used to calculate Composite Selection Signals (CSS) for the comparison of Irish and British breeding Eurasian curlews (*Numenius arquata*) to a population in northern Sweden, using 56 whole-genome sequences.

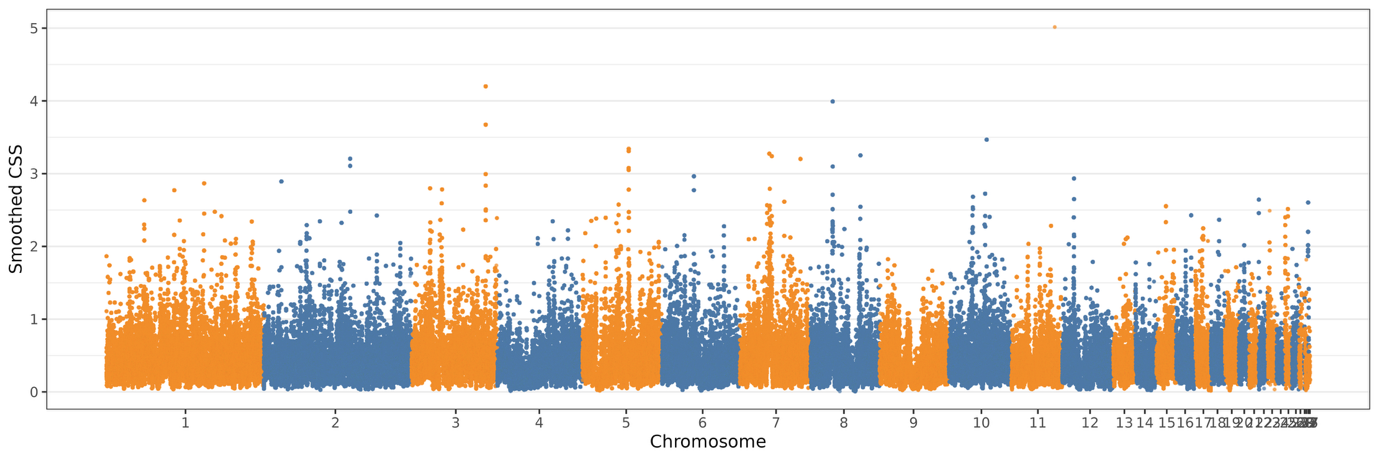

**Fig. S7.** Composite Selection Signal (CSS) scores across the genome for Irish versus British Eurasian curlew (*Numenius arquata*) populations. Peaks indicate regions under putative divergent selection based on combined metrics (*F*_ST_, *XP-EHH*, and *ΔSAF*) smoothed over 20 kb windows.

b

a

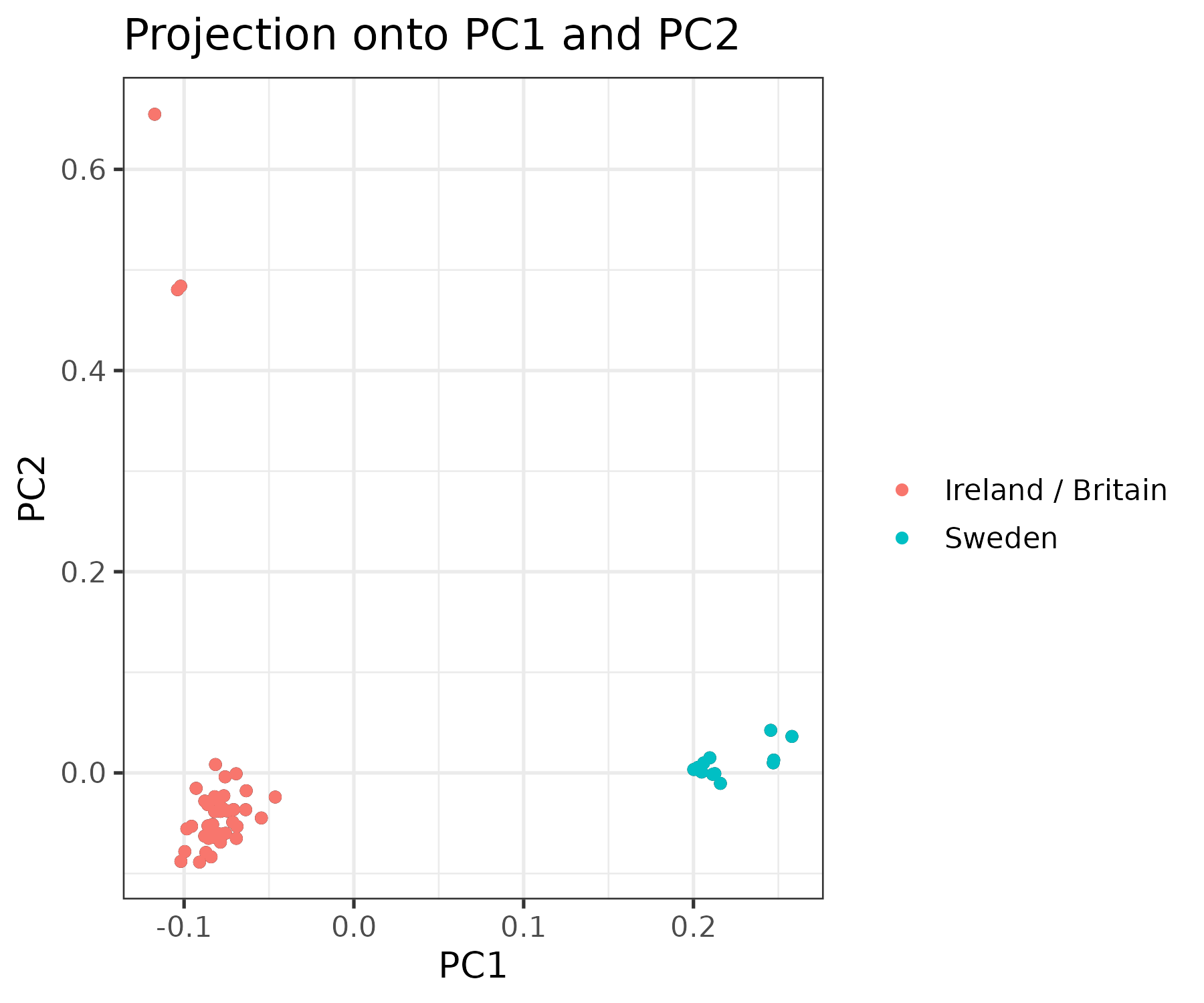

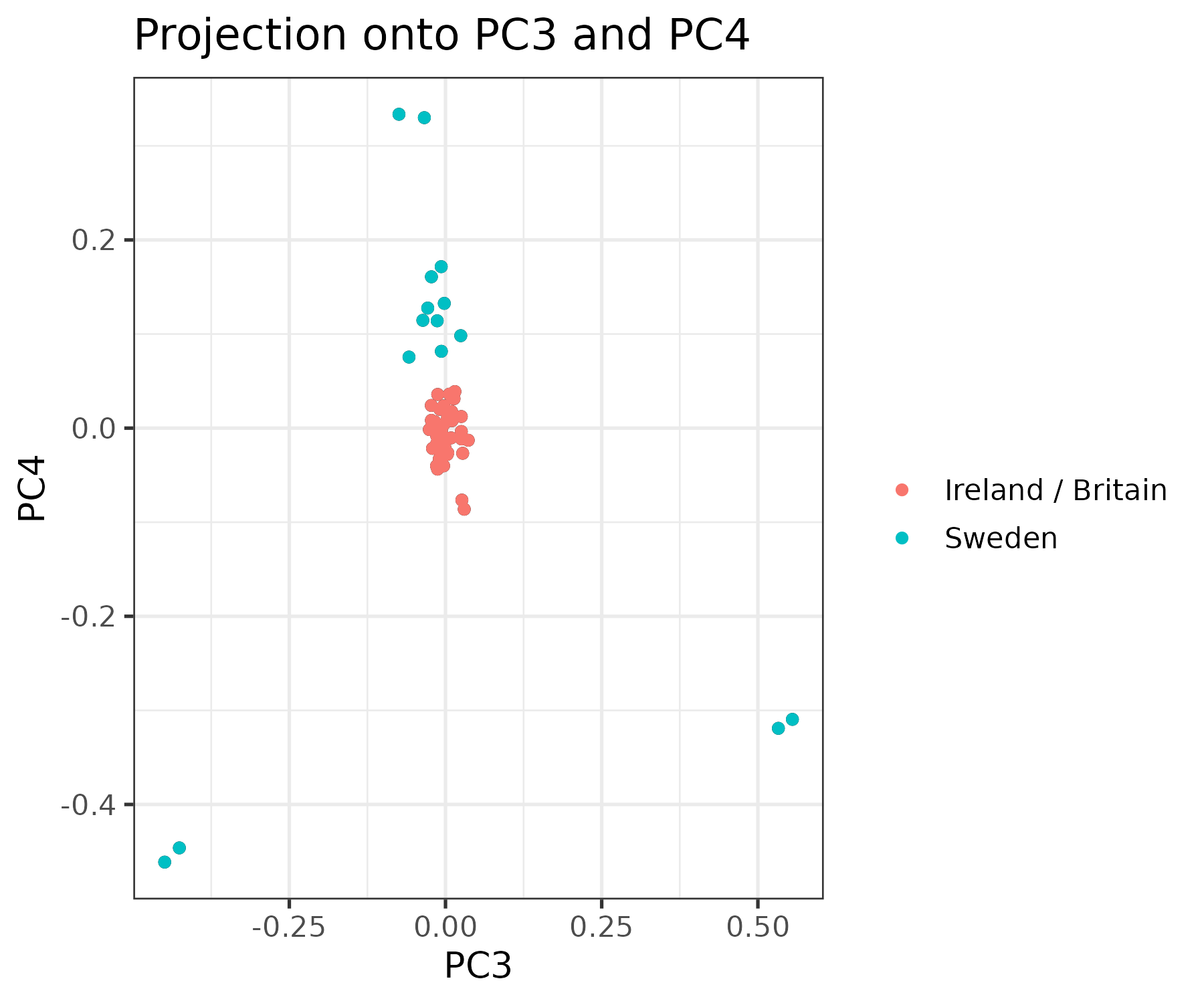

**Fig. S8.** Score plots for a) Principal Components 1 and 2, and b) 3 and 4, which the pcadapt outlier method using 56 Eurasian curlews (*Numenius arquata*) across three study groups in Ireland, Britain, and northern Sweden based on 5,523,293 SNPs

b

a

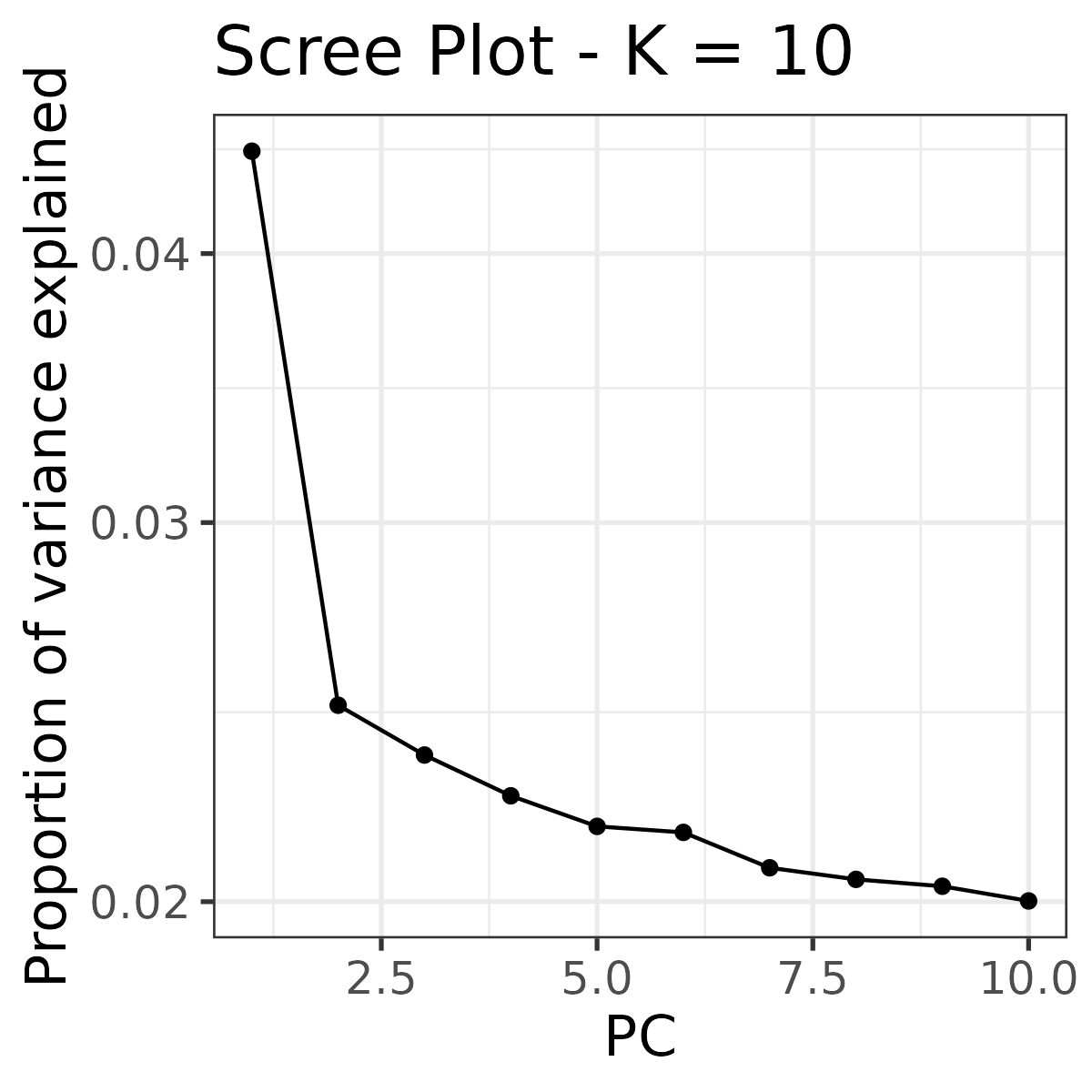

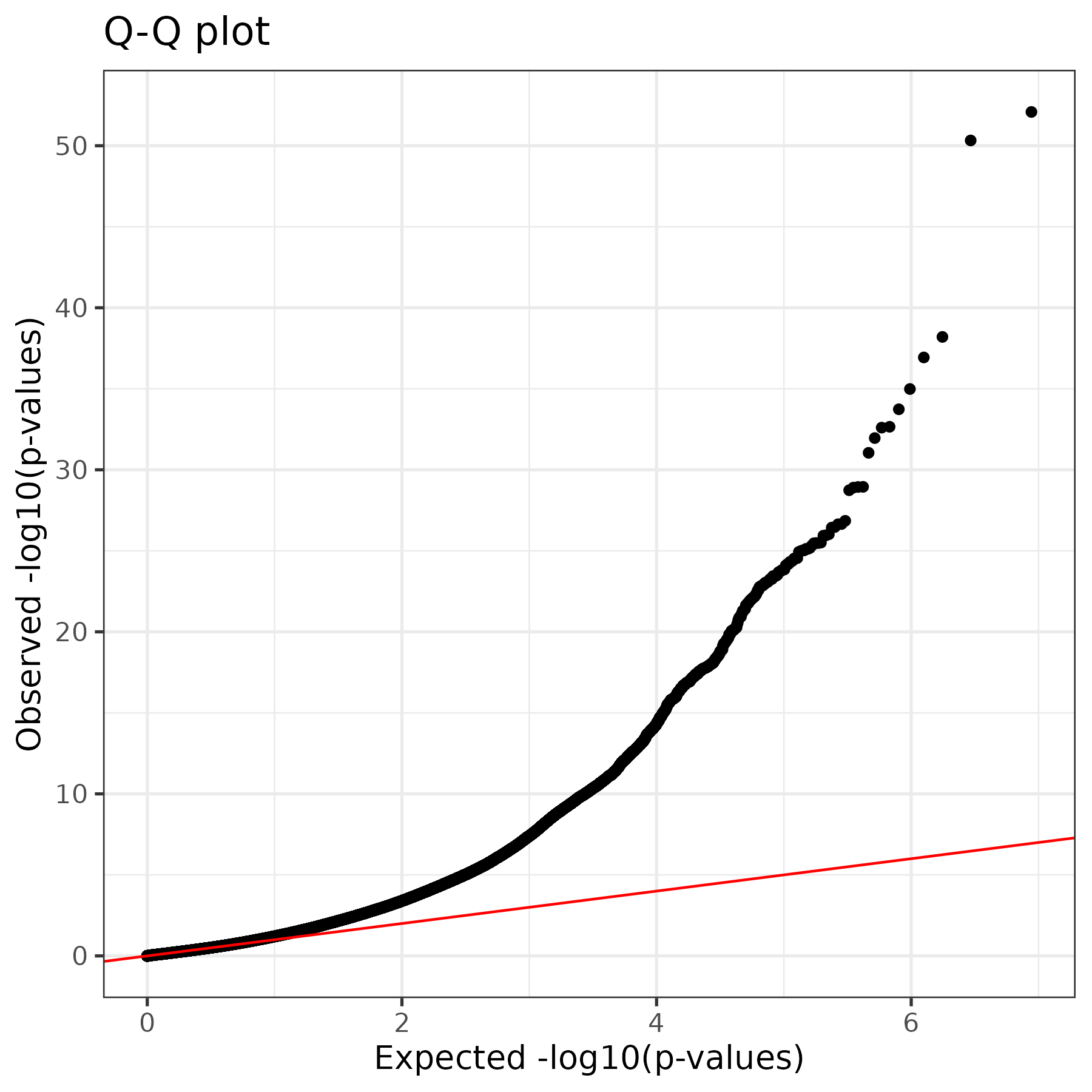

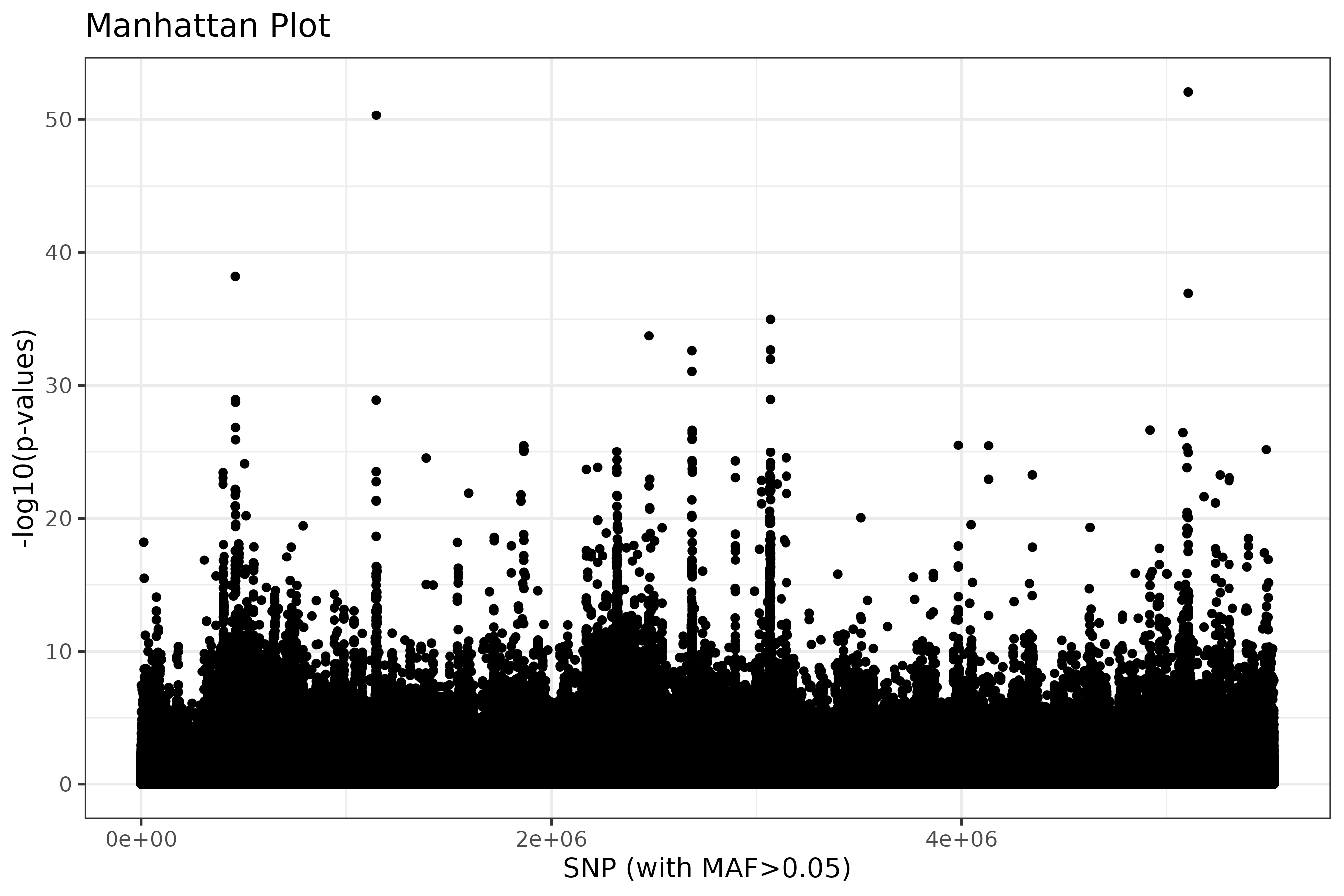

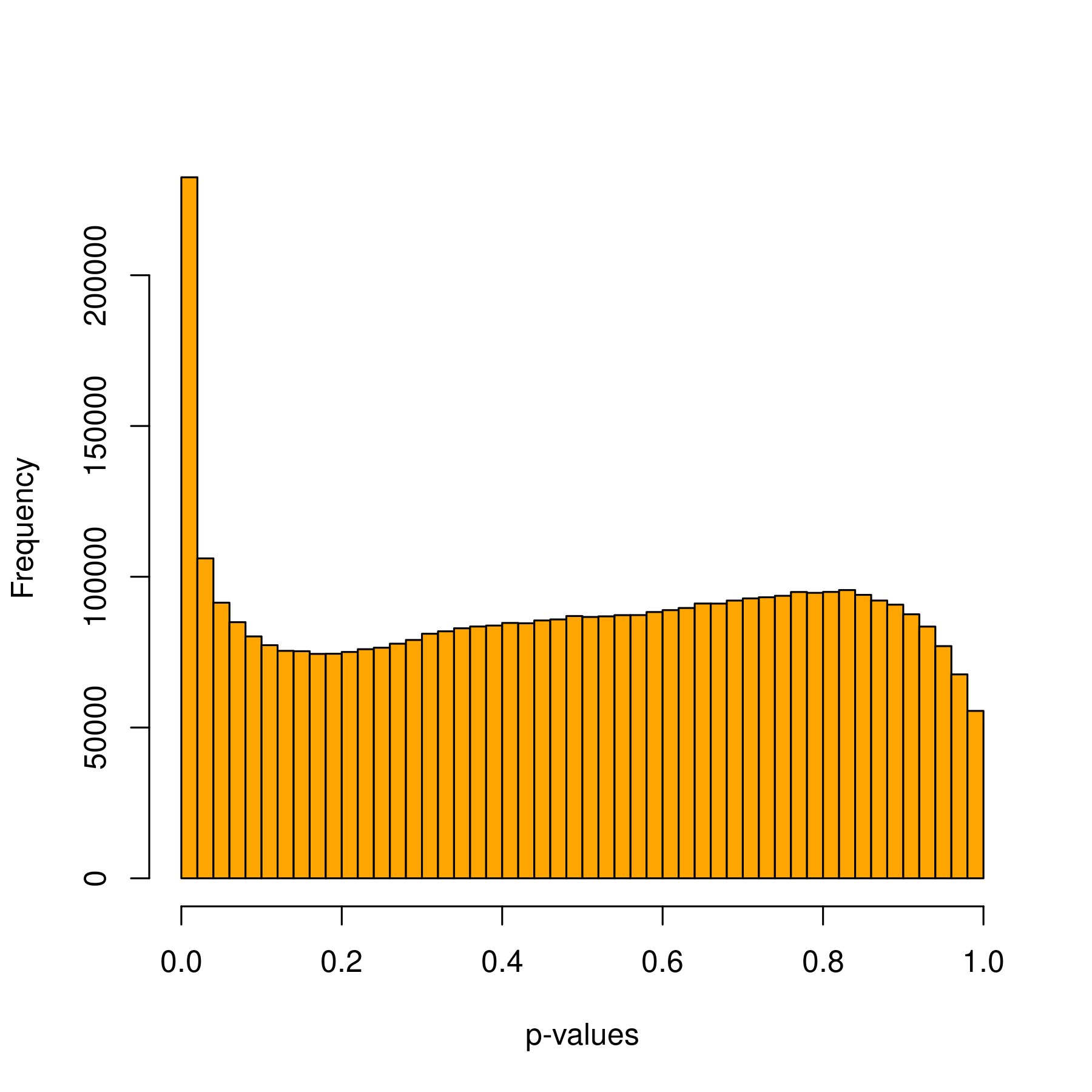

d

c

**Fig. S9.** a) qq plot of −log_10_*P* values for the SNPs, b) histogram of the frequency of *P* values, c) Scree plot produced with pcadapt showing the variance explained by the first 10 principal components and d) Manhattan plot of −log_10_*P* values.

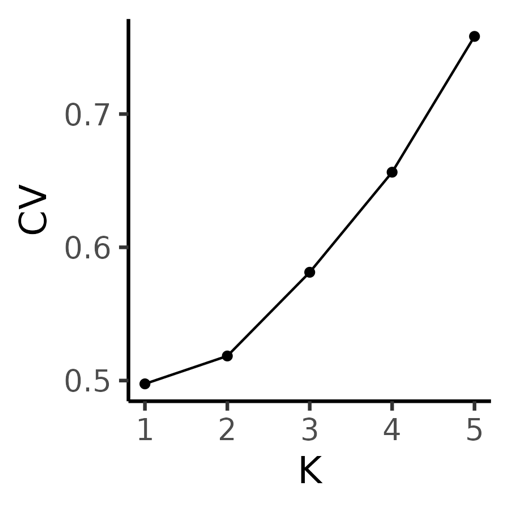

**Fig. S10.** Cross-validation (CV) error from ADMIXTURE analysis for *K* = 1–5, where *K* is the assumed number of ancestral populations. CV error is lowest at *K* = 1 and increases with higher K.

**Table S1.** Summary of whole‑genome resequencing batches for Eurasian curlews (*Numenius arquata*), raw read depth, and post‑mapping coverage. All batches were sequenced on an Illumina NovaSeq X Plus.

| **Batch no.** | **Library Preparation** | **No. sequenced** | **Date** | **Raw depth** | **Post mapping depth** |
| --- | --- | --- | --- | --- | --- |
| P29966 | Illumina DNA PCR-Free Prep (Illumina, Inc., California, USA) | 15 | 30/12/2023 | 20.4x | 14.8x |
| P33209 |  | 18 | 19/11/2024 | 24.9x | 19.4x |
| P34852 |  | 23 | 16/05/2025 | 30.9x | 21.8x |
| 90-1188102853 | NEBNext Ultra II DNA (NEB, Ipswich, MA, USA) | 10 | 18/04/2025 | 33.9x | 21.3x |

**Table S2.** Summary of SNP filtering steps applied to the whole‑genome resequencing dataset of Eurasian curlews (*Numenius arquata*), showing the number of retained variants at each stage.

| **File** | **Filtering step** | **SNPs retained** | **Filtering criteria** | **Analysis** |
| --- | --- | --- | --- | --- |
| 1 | Initial filtering | 15,953,091 | QUAL < 30, MQ < 40.00, SOR > 4.000, QD < 2.00, FS > 60.000, MQRankSum < -12.5, ReadPosRankSum < -8.000 | --- |
| 2 | Only autosomes | 14,934,126 |  | --- |
|  | All sites autosomes | Sites retained: 550,160,281 | QUAL < 30, MQ < 40.00, SOR > 4.000, QD < 2.00, FS > 60.000, MQRankSum < -12.5, ReadPosRankSum < -8.000 | --- |
| 3 | No MAF | 7,513,513 |  | ROH, pixy |
| 4 | MAC < 3 | 5,523,293 | --mac 3  --maxDP 40  --minDP 10  --max-missing 0.9 | Diversity estimates,  Principal component analysis, *F*_ST_ |
| 5 | Linkage disequilibrium | 782,547 | r^2^ = 0.2 | ADMIXTURE |
| 6 | Thinning | 816 – Ireland  818 – Britain  819 – Sweden | --thin 1000000  --max-missing 1 | ONeSAMP |

**Table S3.** Significant Gene Ontology (GO) enrichment terms identified for outlier genes in the Ireland/Britain vs Sweden comparison of the Eurasian curlews (*Numenius arquata*). The table lists enriched Molecular Function (MF) and Cellular Component (CC) categories, including GO term identifiers and adjusted *P*‑values.

| **GO category** | **GO term** | **GO term ID** | ***P*_adj._ value** |
| --- | --- | --- | --- |
| GO:MF | G protein-coupled peptide receptor activity | GO:0008528 | 0.0001 |
| GO:MF | peptide receptor activity | GO:0001653 | 0.0002 |
| GO:MF | peptide binding | GO:0042277 | 0.0343 |
| GO:CC | cytosolic ribosome | GO:0022626 | 0.0084 |
| GO:CC | ribosomal subunit | GO:0044391 | 0.0147 |

**Table S4.** Gene Ontology (GO) enrichment results for the Ireland vs Britain comparison of the Eurasian curlews (*Numenius arquata*). The table lists enriched Molecular Function (MF) and Cellular Component (CC) categories, including GO term identifiers and adjusted *P*‑values based on Composite Selection Signals (CSS).

| **GO category** | **GO term** | **GO term ID** | ***P*_adj._ value** |
| --- | --- | --- | --- |
| GO:MF | volume-sensitive anion channel activity | GO:0005225 | 0.0500 |
| GO:BP | positive regulation of vascular endothelial cell proliferation | GO:1905564 | 0.0466 |
| GO:CC | DNA-dependent protein kinase complex | GO:0070418 | 0.0072 |
| GO:CC | intracellular anatomical structure | GO:0005622 | 0.0170 |
| GO:CC | cytosol | GO:0005829 | 0.0340 |
| GO:CC | Ku70:Ku80 complex | GO:0043564 | 0.0499 |

**Appendix 2.** List of outlier genes under putative divergent selection identified through Composite Selection Signal (CSS) analysis between a population of Eurasian curlews (*Numenius arquata*) in Ireland and Britain, contrasted with a Swedish population. (Supplementary_Table_2.xlsx)

**Appendix 3.** List of outlier genes under putative divergent selection identified through principal components (R package: pcadapt) between a population of Eurasian curlews (*Numenius arquata*) in Ireland and Britain, contrasted with a Swedish population. (Supplementary_Table_3.xlsx)

Supplementary Methods

To calculate Composite Selection Signals (CSS), the phased SNP dataset was used. The CSS statistic (Randhawa et al. 2014) can be used to detect regions or genes under divergent selection by combining multiple statistics into one measure. Three metrics were calculated to compare the combined Irish/British breeding population samples with the Swedish samples. Firstly, SNP-by-SNP *F*_ST_ (Weir and Cockerham 1984) was calculated with VCFtools. Second, *XP*-*EHH* (cross‑population extended haplotype homozygosity) statistics were calculated in selscan v1.2.0 (Rahman et al. 2025), using default parameters. The raw *XP*-*EHH* scores were normalised using the --*norm* function in selscan. Third, the directional change in the selected allele frequency (*ΔSAF*) statistic was calculated using the --*freq* function in VCFtools to output allele frequencies, and a custom R script to calculate *ΔSAF* and standardise to *Z* ~ *N*(0,1) (Randhawa et al. 2014).

The *F*_ST_, *XP*-*EHH*, and *ΔSAF* statistics were joined by genomic location (chromosome:position) and combined into a CSS statistic for each SNP using a custom R script following (Randhawa et al. 2014). Briefly, each statistic was ranked across all SNPs and converted to fractional ranks. These were then converted to z-scores, averaged, and converted to *P*-values and the CSS were defined as −log_10_(*P*-value). Each SNP CSS score was then averaged over a 20kb window to reduce statistical noise. To identify outlier SNPs, a significance threshold was set to identify SNPs in the top 0.1% of CSS scores. These SNPs then had to be flanked (± 0.5 Mb) by at least five other SNPs in the top 1% of CSS values (Han et al. 2023; Randhawa et al. 2014; Ward et al. 2024). Regions were then defined as the span between the outermost flanking SNPs in the top 1%. If these regions were within 250 kb of each other, they were merged. Genes within these regions, or ±200 kb around them, were extracted from the genome annotation.

Supplementary Discussion

Energetics, appetite, and circadian regulation

Swedish curlews may be under higher energetic demands than Irish and British birds due to a shorter breeding season (Pederson et al. 2022) and higher migration burden (Brown 2015). Most work on candidate genes and migration concerns passerines (Gu et al. 2024), and there is no consistent pattern of selection on specific genes or gene regulatory elements (GREs) in birds (Delmore et al. 2020; Gu et al. 2024; Lugo Ramos et al. 2017; Sokolovskis et al. 2023). Pre-fuelling timing influences migration, and the associated increase in appetite may involve circadian regulators. We identified *ID2*, a circadian depressor (Ward et al. 2010) associated with feed efficiency in chickens (Shah et al. 2019). The *MC3R* gene was also identified as an outlier, which is involved in appetite regulation. The most significantly overrepresented GO term was G protein coupled peptide receptor (GPCR) activity, with 14 genes, including *MC3R* and *CCKAR*, both of which have roles in avian appetite (Aderibigbe et al. 2022; Li et al. 2025). The full list of the 14 GPCR genes is *CCKAR, NMBR, CYSLTR2, EDNRB, MLNR, GAL, MC3R, OXTR, MAS1, NPY4R, AVPR1B, OPRK1, NPBWR1,* and *P2RY8*. Downregulation of *NPY4R* is linked to appetite suppression (Ghashghaei et al. 2026), while *GAL* is associated with increased feeding (Tachibana et al. 2008) and is disrupted by heat stress (Uyanga et al. 2023). Circadian and metabolic pathways are interconnected, and dysregulation of genes encoding circadian regulators such as *ID2* could alter appetite and energy balance. Therefore, these genes collectively represent putative targets of selection related to energetic demands of a shorter breeding season or pre‑migratory fuelling; however, functional validation in shorebirds is lacking. Six of the genes identified here as outliers were significantly differentially expressed between migratory and non-migratory Swainson's thrush (Louder et al. 2024). These were *FAM167A, ID2*, *PER3, PPP1R3C, RASSF7*, and *TES*.

Photoperiod is another potential driver of divergence between these populations and is related to circadian rhythms. Genes identified as involved in energetics or migration that could also play a role in this include *MC3R*, *ID2* and *PER3*.

Olfaction has several functions in birds including foraging, individual recognition, partner selection, predator detection and nesting (Creece et al. 2025). Some research has implicated olfactory cues in bird movement including long-distance foraging in seabirds (Abolaffio et al. 2018) and homing in pigeons (Bonadonna and Gagliardo 2021). The *TAAR5* olfactory gene (Manzini et al. 2022) is expressed in the chicken olfactory epithelium along with *GNAL,* which encodes a component of the olfactory signalling pathway. Two olfactory receptor genes (*OR4A15, OR52D1*) were also identified in the outlier analysis between Irish/British and Swedish curlew, although no work on bird migration has been done with these particular genes. Other potential candidate genes include *PIEZO2*, which encodes a mechanosensory ion channel protein involved in tactile feeding behaviour and mechanosensory function (Schneider et al. 2019), and is also implicated in mammalian proprioception (Woo et al. 2015).

Immune genes

Variation at TLR gene loci has been linked to reduced survival in the pale-headed brushfinch (*Atlapetes pallidiceps*), likely as an adaptation to localised environmental conditions in a small population (Hartmann et al. 2014). In the endangered Stewart Island robin (*Petroica australis rakiura*) (Grueber et al. 2013) and Attwater's prairie-chicken (*Tympanuchus cupido attwateri*) (Bateson et al. 2016), specific TLR locus genotypes have been associated with survival.

The *CX3CR1* gene, identified as an outlier, encodes a receptor for *CX3CL1*, which is involved in the immune response in chicken macrophages and T cells (Vu et al. 2025). In addition, *XCR1* is highly expressed in chicken spleen, and is involved in the innate immune responses, most notably, it is expressed on cross-presenting dendritic cells (DCs) to counter viruses and other intracellular pathogens (Wu et al. 2023).

Several cytokine and immune-signalling-related genes also appeared as outliers. The *IL19* gene encodes the IL-19 cytokine, which promotes the expression of anti- and pro-inflammatory cytokines in chicken immune cells (Kim et al. 2009). The *TNFAIP1* gene is inherently more highly expressed in IBV-resistant compared with IBV-susceptible chicken lines (Smith et al. 2015). The *GAB3* gene is significantly upregulated in erythroid chicken cells during Avian Pathogenic *Escherichia coli* (APEC) infection (Cai et al. 2026) and, in mammals, is required for IL-2/IL-15-mediated NK cell activation, which contributes to innate antiviral and anti-tumour responses (Sliz et al. 2019). Additional outlier genes with potential immune functions included the proteinase inhibitor-encoding *SERPINA3*, which was upregulated in chicken liver and serum during bacterial infection (Polansky et al. 2018). Several additional outlier genes were associated with broader immune‑related GO categories, including *immune system process* (*HMGB3*), *innate immune response* (*HMGB3, RB1CC1*, and *RNF7*), and *inflammatory response* (*PARK7, AGR2, FAM210B, TAC1*, and *TP73),* with *OPRK1* appearing under the general *immune response* term.

Thermal adaptation

A study that exposed tree swallow (*Tachycineta bicolor*) nestlings to a temperature rise of +4.5°C showed significant differential gene expression in peripheral blood for several genes we also identified, including *MITD1*, *TLR2*, *SMARCA1,* and *PCMTD1* (Woodruff et al. 2025). Other outliers potentially involved in heat include *PSMD10*, which encodes a chaperone protein involved with the assembly of the 26S proteasome, which is activated under heat stress in mammals (Lee and Goldberg 2022) and downregulated in heat-stressed chickens (Wang et al. 2014); *DUSP1* (heat-responsive in chickens) (Wang et al. 2025); *AQP11,* which is significantly upregulated by heat stress in chickens (Aloui et al. 2024); and *TLR2,* which shows increased expression in chickens under heat stress (Abdelaziz et al. 2024). In addition, domestic turkeys (*Meleagris gallopavo*) cold-treated for three days showed differential expression for several of our outlier genes, including *KLHL38, PSMD10,* and *RB1CC1* (Reed et al. 2025). This study, along with multiple other scientific findings, suggests that thermal stress affects the expression of some of the outlier genes identified in this study. However, in the absence of thermoregulatory pathway overrepresentation and with many of the genes being general stress-, metabolic-, and immune-related, this suggests that thermal selection is unlikely to be a major driver of divergence between the populations.

Embryogenesis and development

Genes with potential roles in embryogenesis include: *BMP4*, *GSC*, *HOXA7*, *HOXA9*, *HOXA11*, *MYF5*, several fibroblast growth factor (FGF) genes (involved in cell growth, tissue repair, and embryonic development), *SALL4* and *NEUROG1*. HOX genes have conserved roles in axial patterning and vertebrate gastrulation (Davis et al. 1995; Fromental-Ramain et al. 1996; Wang et al. 2026), but some outliers typically associated with embryogenesis also have other roles. The *BMP4* gene is involved in several aspects of vertebrate development, including gut nervous system development (Kovács et al. 2023), limb development (Pizette and Niswander 1999) and fat deposition (Cheng et al. 2016). The *GSC* gene is expressed very early in embryo development and organisation (Izpisúa-Belmonte et al. 1993). Several HOX genes were detected as outliers, including *HOXA7, HOXA9,* and *HOXA11*, which have conserved roles in axial patterning and vertebrate gastrulation (Davis et al. 1995; Fromental-Ramain et al. 1996; Moreau et al. 2019; Wang et al. 2026). These genes were identified with pcadapt and occur in a cluster on curlew chromosome 7 (NC_133582), so it is difficult to determine the specific HOX genes under selection. The *MYF5* gene is involved in early muscle development and birth weight (Genxi et al. 2014) and *FGF3, FGF6, FGF13,* and *FGFBP2* are fibroblast growth factors with well-documented embryonic roles. In this regard, the *FGF3* gene is involved in cochlear development (Olaya-Sánchez et al. 2017) and the hypothalamo-neurohypophyseal axis (Liu et al. 2013). *FGF6* is involved in myogenesis (Smith and Jerome-Majewska 2024), *FGF13* is expressed in chicken limbs during development (Munoz-Sanjuan et al. 2000), and *FGFBP2* is highly expressed in tibial development (Lu et al. 2024). Other genes involved in early development include *NEUROG1* (Bina et al. 2023), *FLRT3* (Tomás et al. 2011), *WNT8B* (Garda et al. 2002), *MAB21L1* (Dai et al. 2014), and *DRAXIN* (Islam et al. 2009). Additional genes with roles in embryogenesis include *TCF21, NELL1, SALL4, TP73,* and *RB1CC1.*
